## Supplemental Material for "Copy number footprints of platinum-based anticancer therapies"

### Supplementary Figures

#### Figure S1

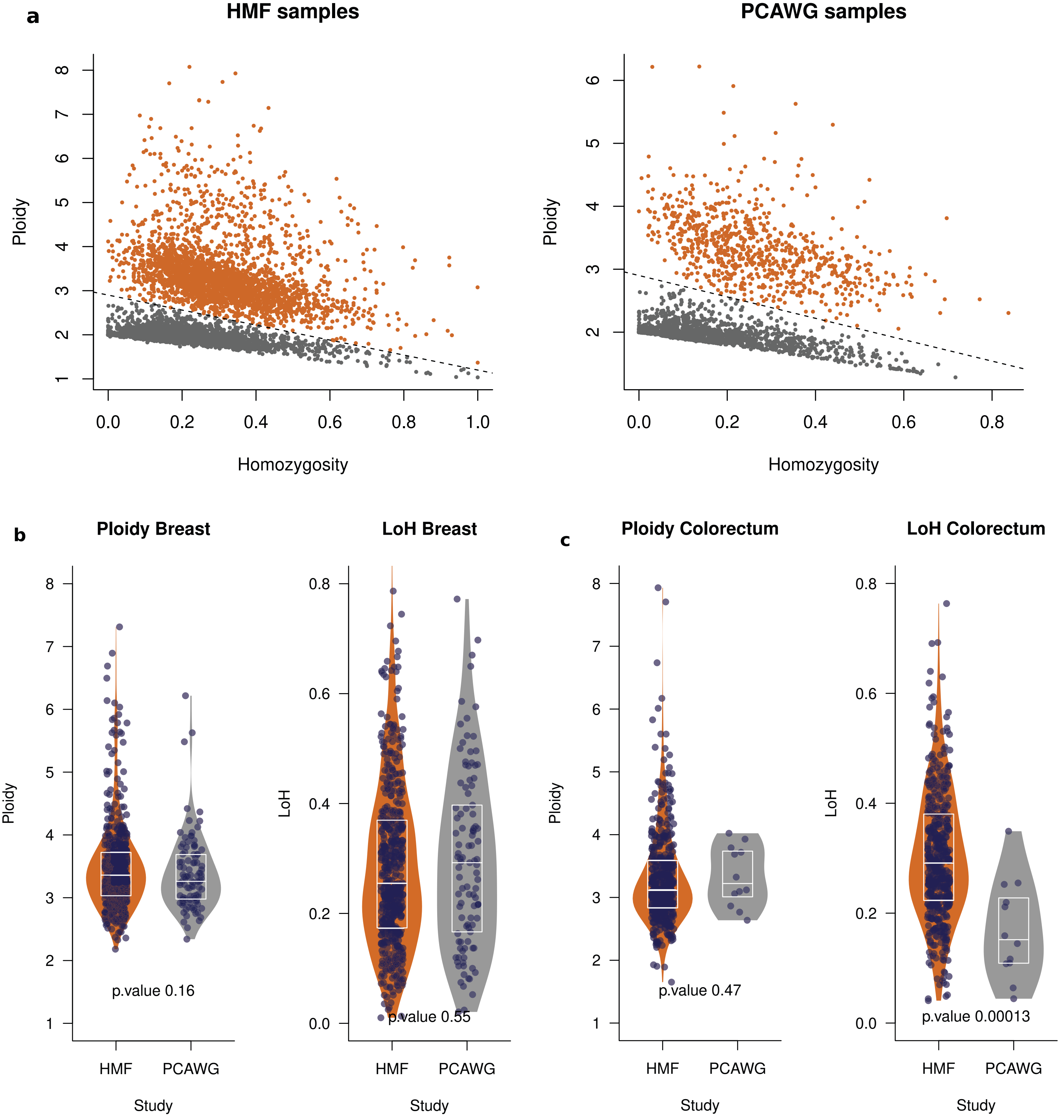

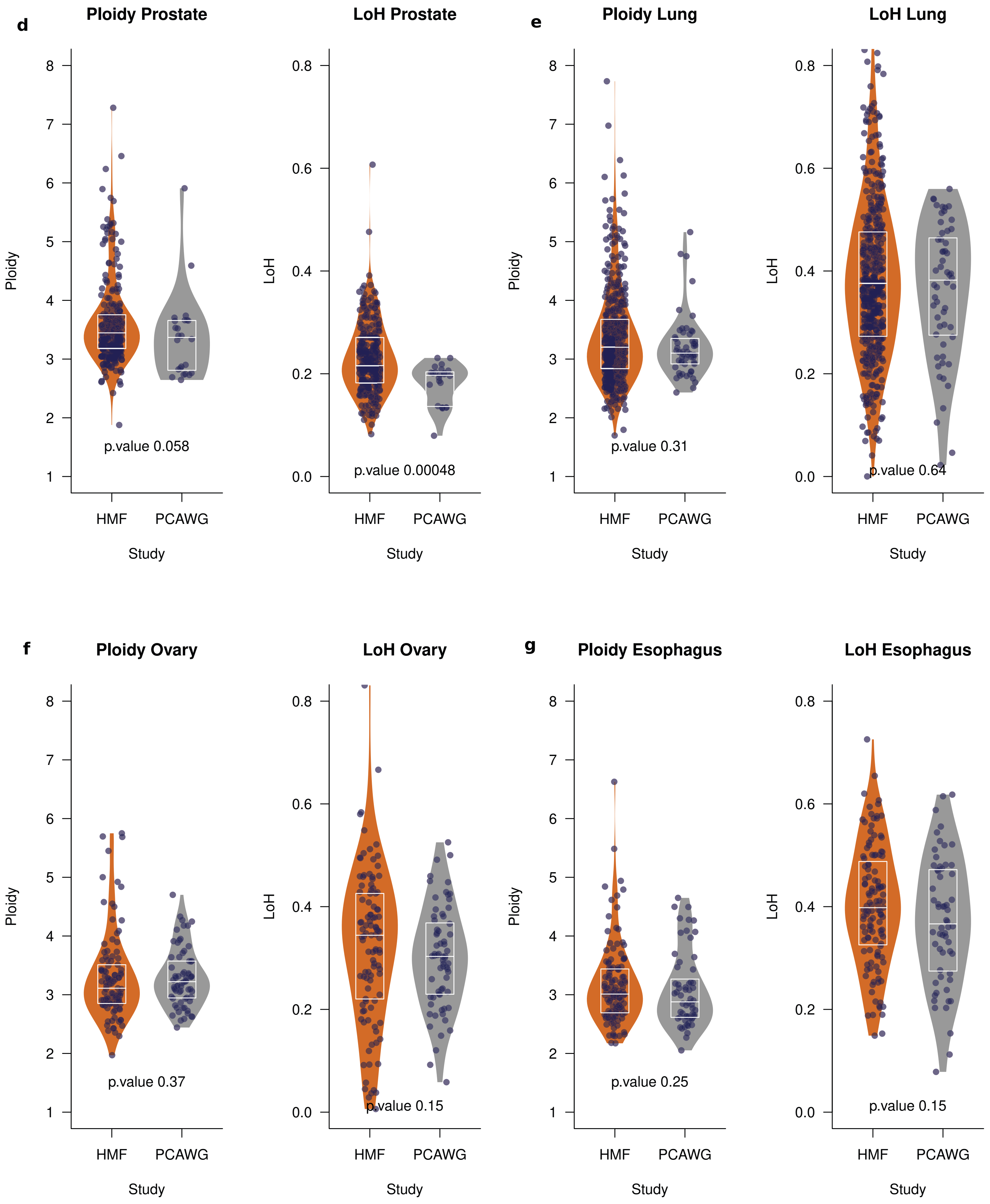
d
Ploidy Prostate
LoH Prostate
e
Ploidy Lung
LoH Lung
Ploidy
LoH
p.value 0.058
p.value 0.00048
p.value 0.31
p.value 0.64
HMF
PCAWG
Study
f
Ploidy Ovary
LoH Ovary
g
Ploidy Esophagus
LoH Esophagus
p.value 0.37
p.value 0.15
p.value 0.25
p.value 0.15

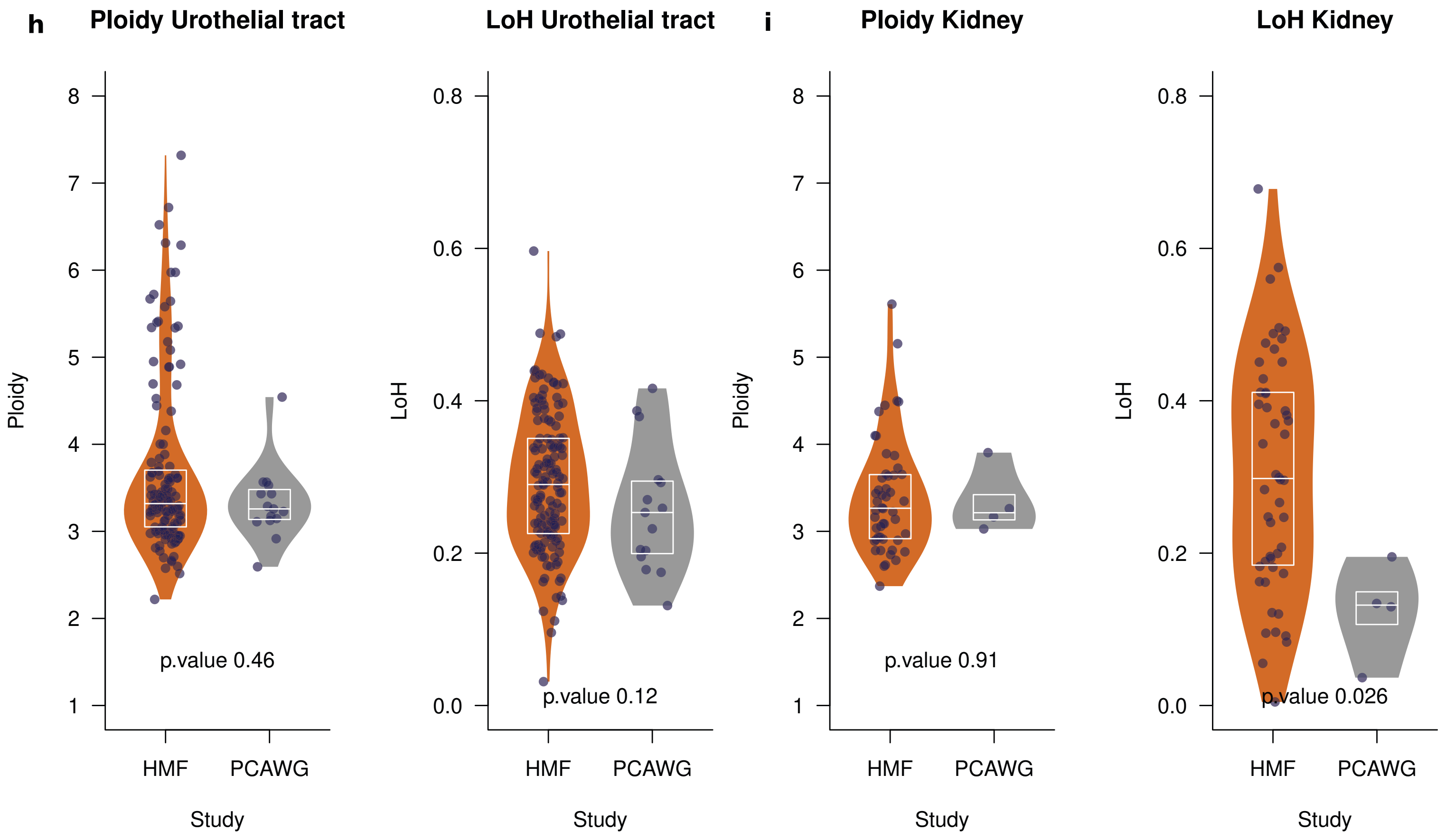


**Figure S1. Primary and metastatic WGD tumors**

a) Separation between WGD and non WGD tumors across the primary (PCAWG) and metastatic (HMF) cohorts on the basis of their ploidy and level of homozygosity. Similar to Figure 1a, but with the tumors of the two cohorts in separate panels.

b-i) Distribution of ploidy (left) and fraction of the genome with LOH (right) of WGD tumors of different cancer types across primary and metastatic cohorts. Similar to Figure 1c, but separated by cancer types represented in both, PCAWG and HMF cohorts. The boxes inside the violin plots delimit the first, second and third quartiles of the distribution. All tumors in each group are represented as dots. P-values were derived from a two-tailed Wilcoxon-Mann-Whitney test.

#### Figure S2

**WGD Taxane**
**Breast_ER–positive/HER2–negative**

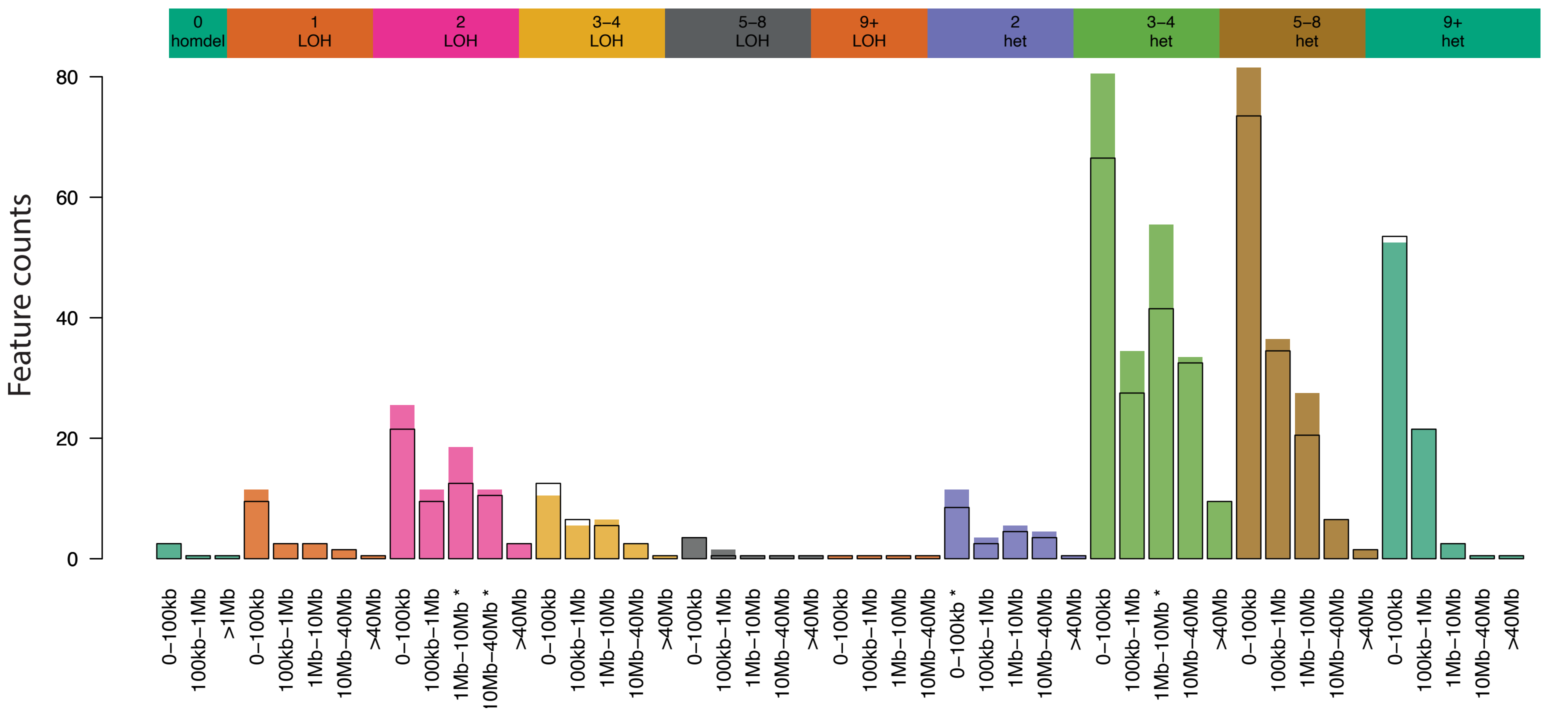


**WGD Taxane**
**Breast_Triple negative**

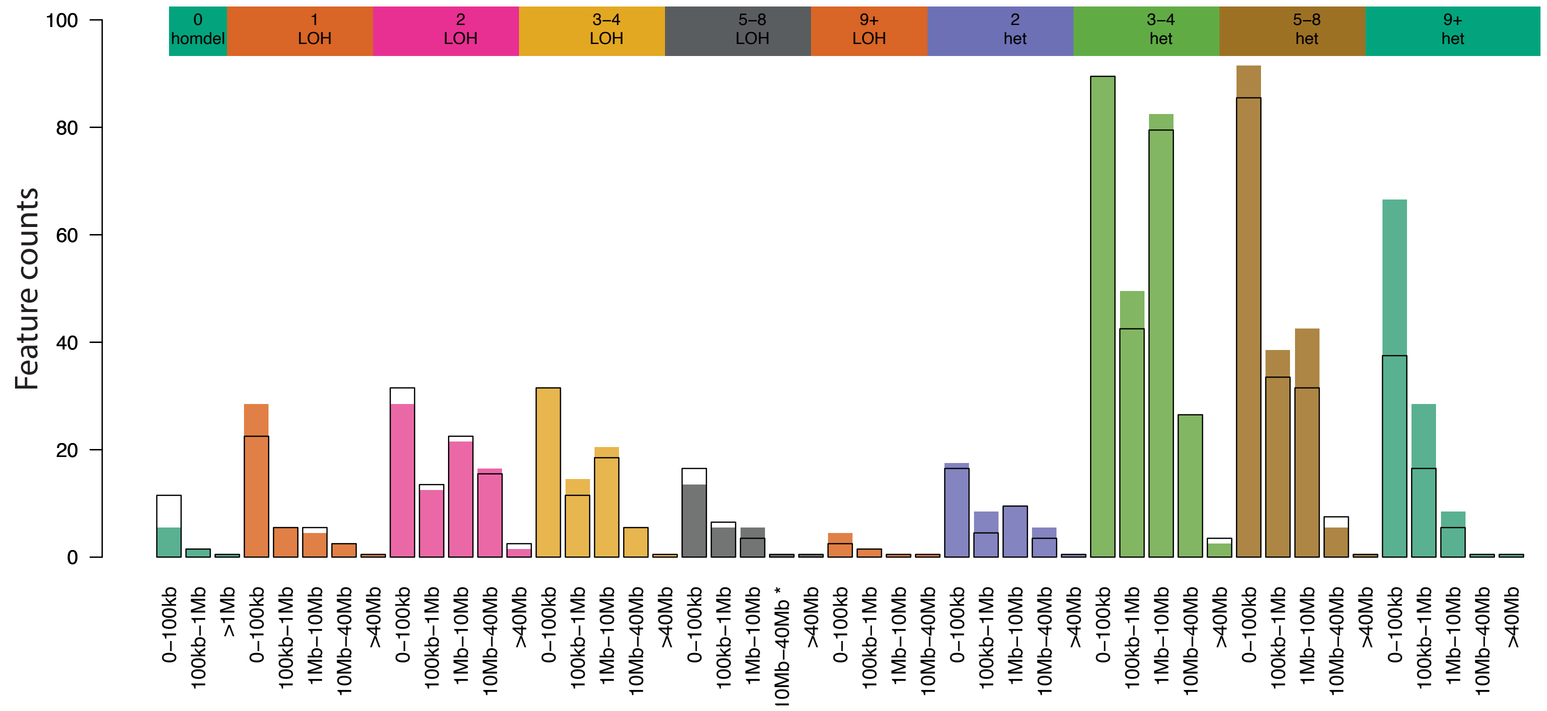

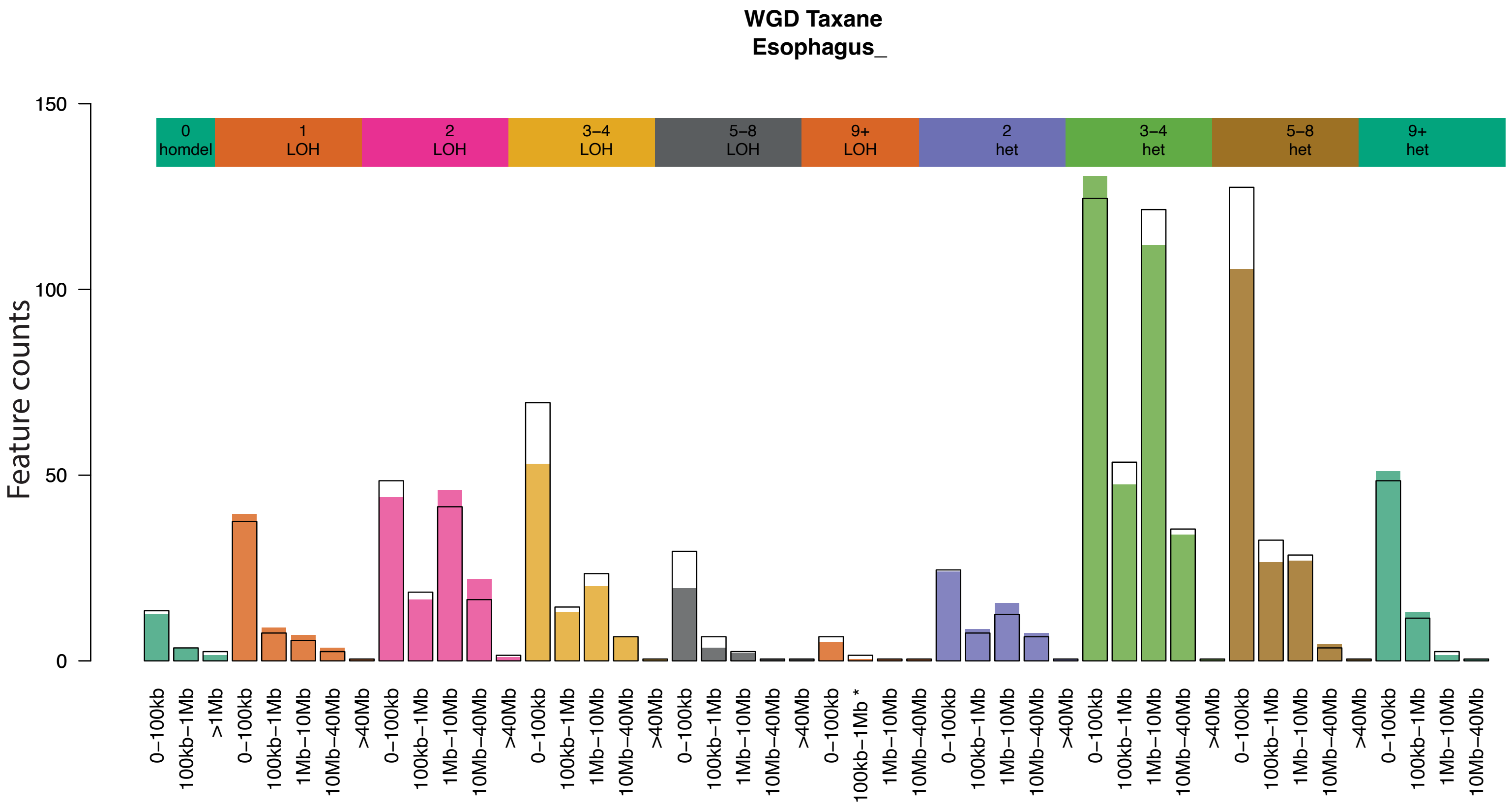
WGD Taxane
Esophagus_
Feature counts
0 homdel
1 LOH
2 LOH
3–4 LOH
5–8 LOH
9+ LOH
2 het
3–4 het
5–8 het
9+ het
0–100kb
100kb–1Mb
>1Mb
1Mb–10Mb
10Mb–40Mb
>40Mb
100kb–1Mb *

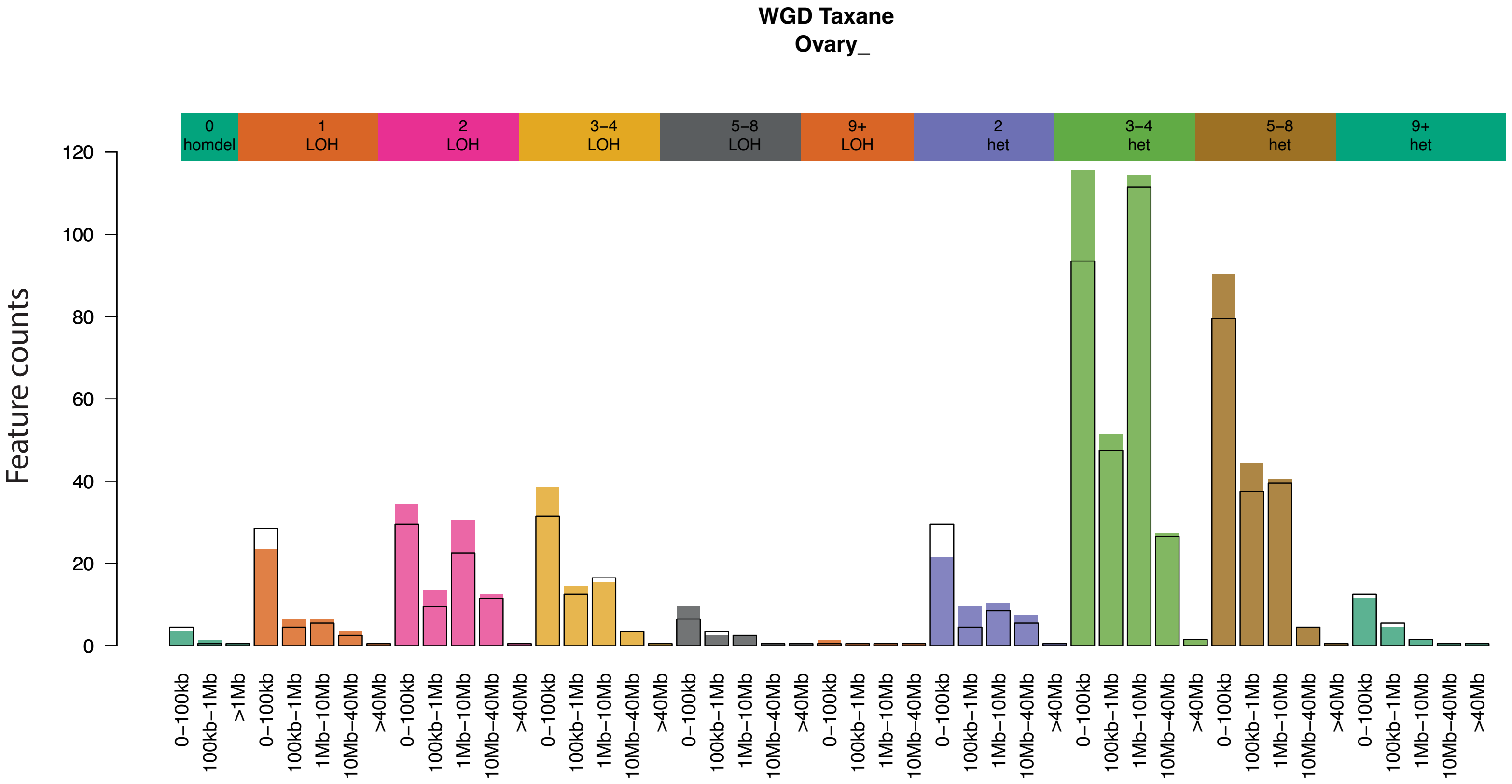
WGD Taxane
Ovary_
Feature counts
0 homdel
1 LOH
2 LOH
3–4 LOH
5–8 LOH
9+ LOH
2 het
3–4 het
5–8 het
9+ het
0–100kb
100kb–1Mb
>1Mb
1Mb–10Mb
10Mb–40Mb
>40Mb

**Figure S2.** Results of the comparison of CN categories across WGD ovarian, esophageal, breast triple negative and breast ER+/HER2- tumors exposed or unexposed to taxanes in the HMF cohort. Significant comparisons (uncorrected p-value below 0.05) are highlighted with asterisks below each bar.

**Figure S3**

**a**

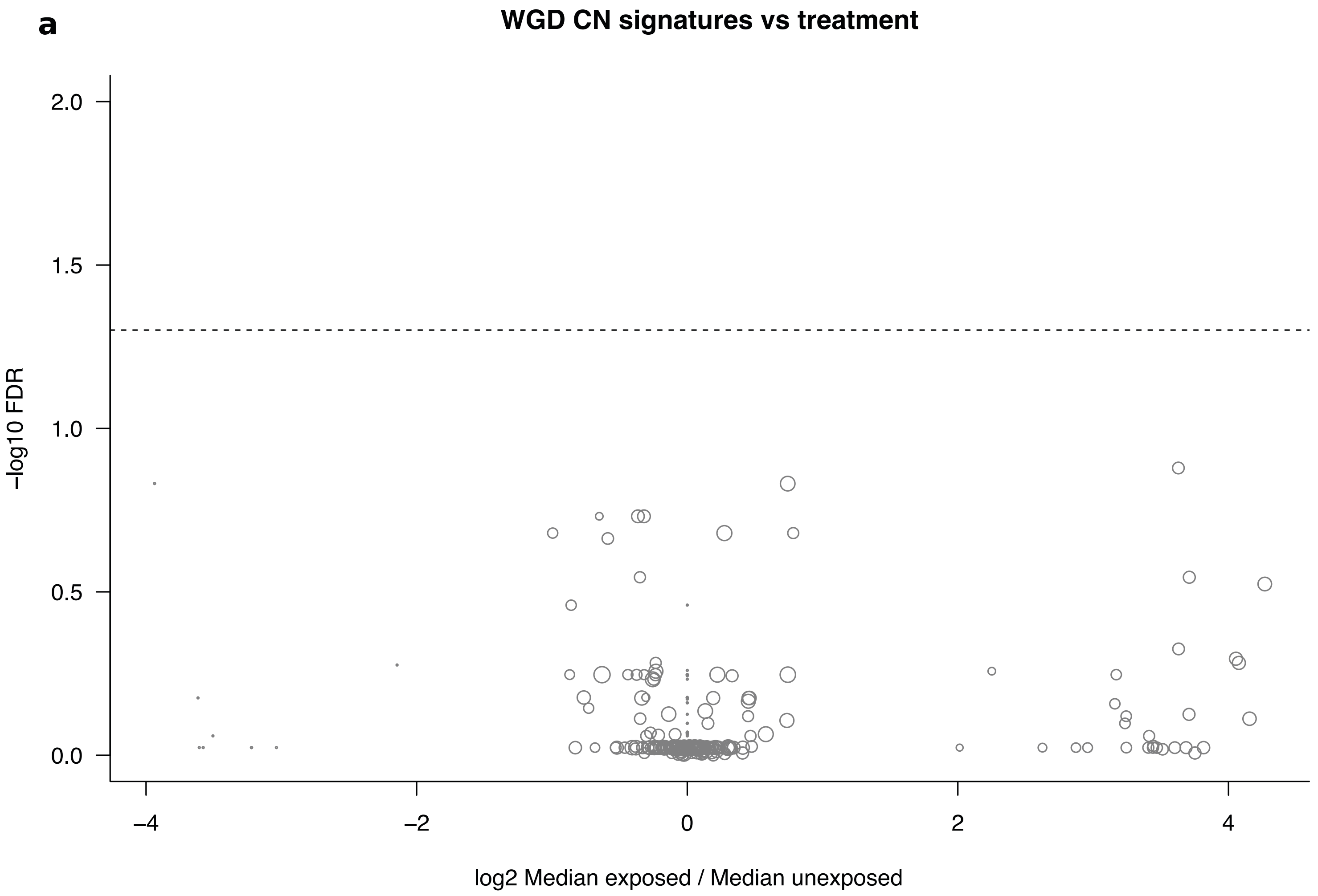

WGD CN signatures vs treatment
2.0
1.5
1.0
0.5
0.0
−log10 FDR
−4
−2
0
2
4
log2 Median exposed / Median unexposed

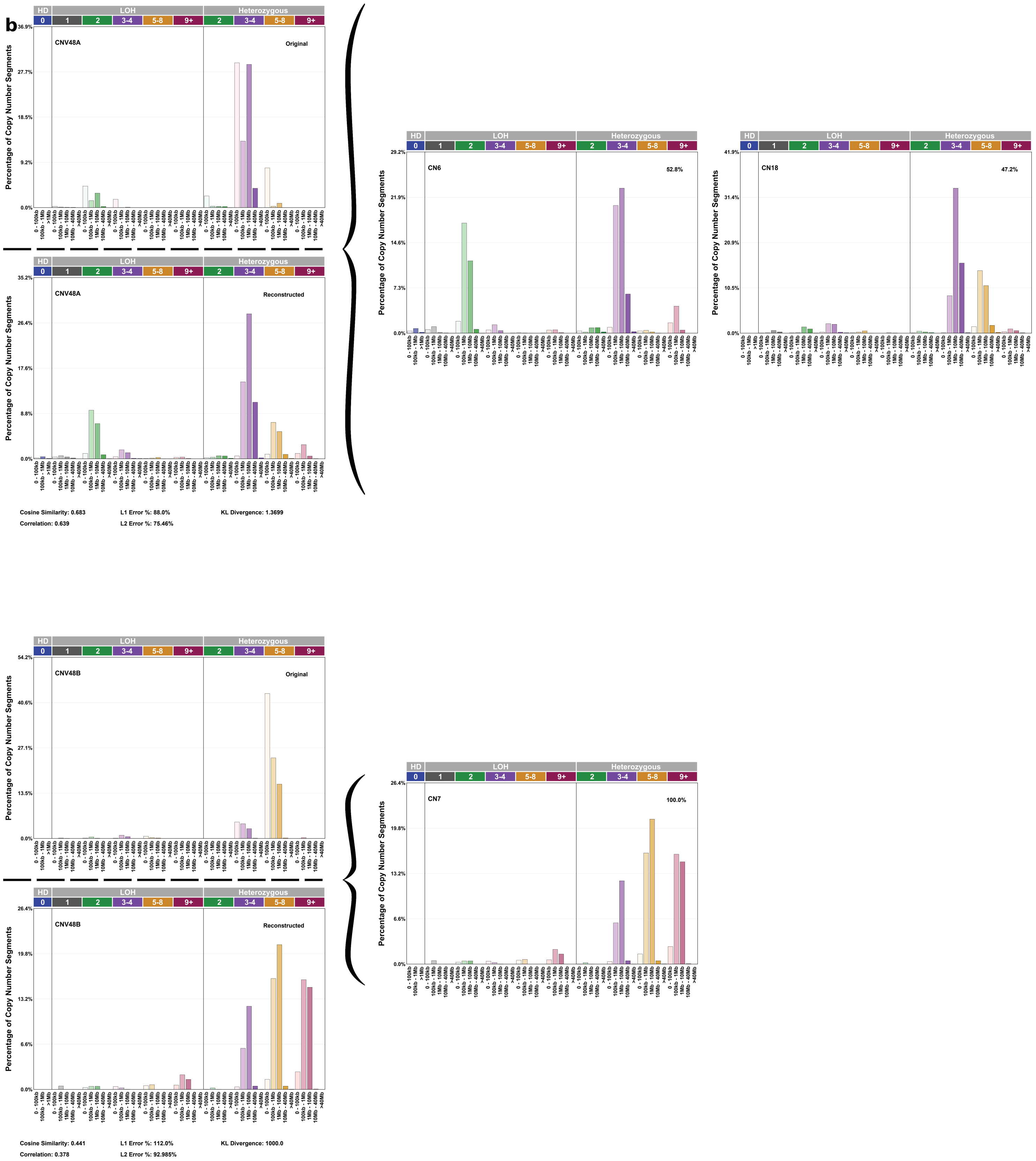
b
HD
LOH
Heterozygous
CNV48A
Original
Reconstructed
Percentage of Copy Number Segments
Cosine Similarity: 0.683
L1 Error %: 88.0%
KL Divergence: 1.3699
Correlation: 0.639
L2 Error %: 75.46%
CN6
52.8%
CN18
47.2%
CNV48B
Original
Reconstructed
Cosine Similarity: 0.441
L1 Error %: 112.0%
KL Divergence: 1000.0
Correlation: 0.378
L2 Error %: 92.985%
CN7
100.0%

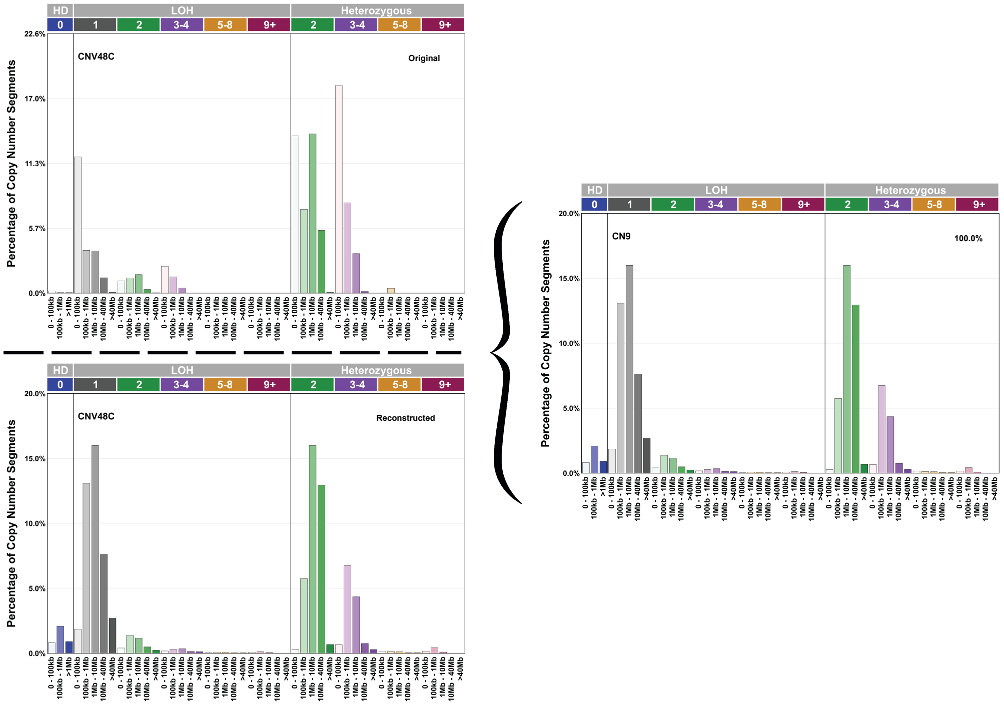


Cosine Similarity: 0.544
Correlation: 0.429
L1 Error %: 97.0%
L2 Error %: 95.759%
KL Divergence: 1.2671

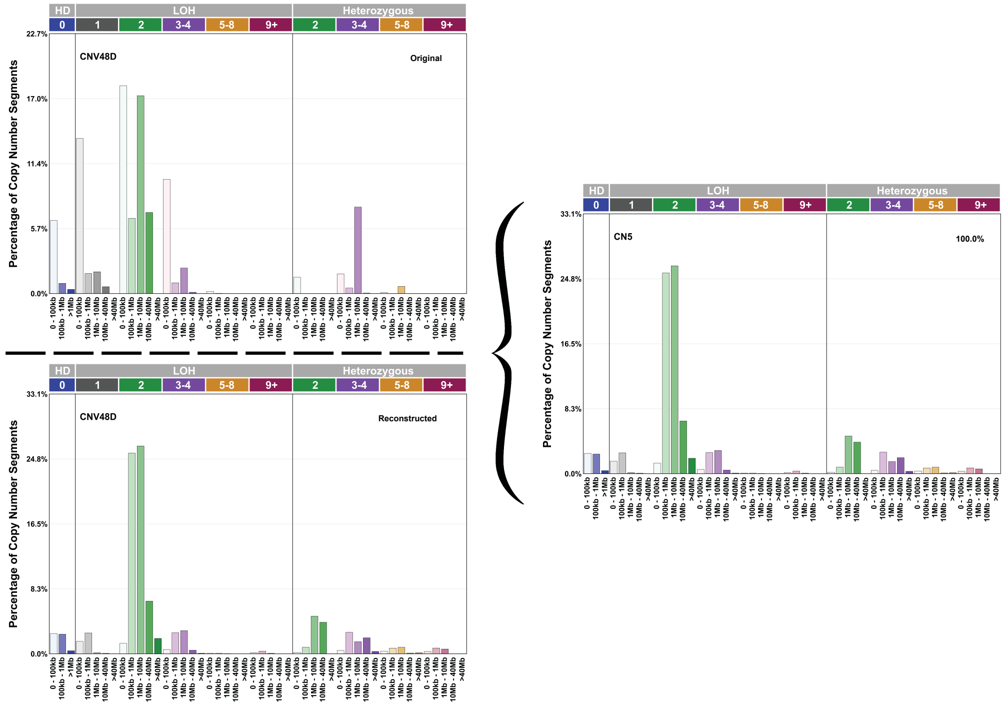


Cosine Similarity: 0.595
Correlation: 0.518
L1 Error %: 107.0%
L2 Error %: 98.471%
KL Divergence: 1.1467

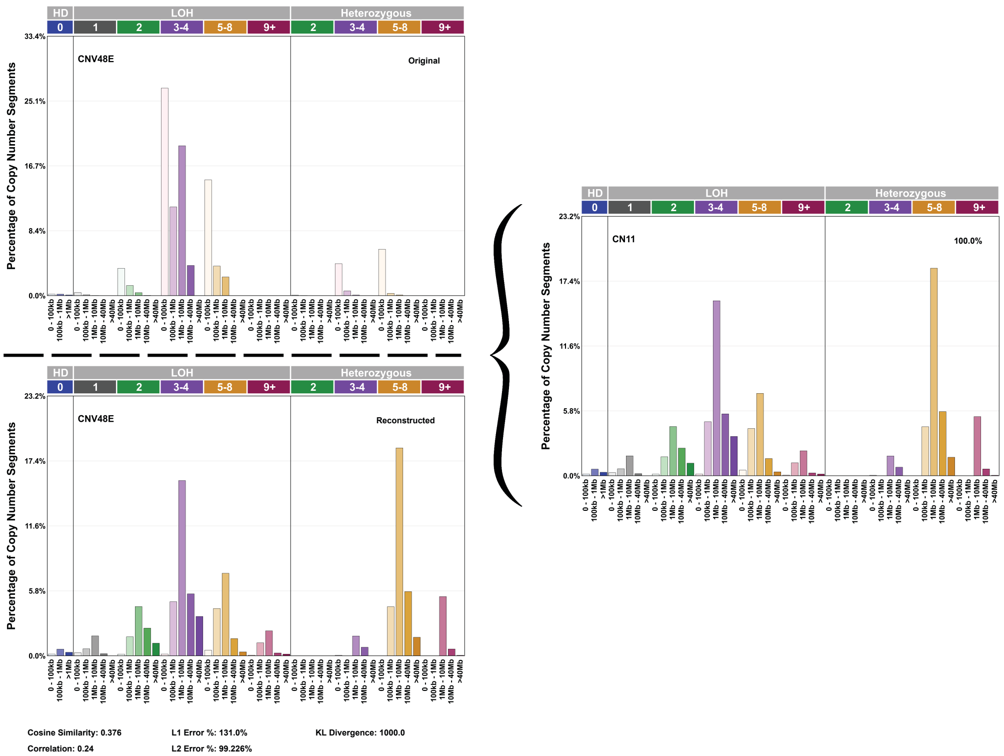
HD
LOH
Heterozygous
CNV48E
Original
Reconstructed
CN11
100.0%
Percentage of Copy Number Segments
Cosine Similarity: 0.376
Correlation: 0.24
L1 Error %: 131.0%
L2 Error %: 99.226%
KL Divergence: 1000.0

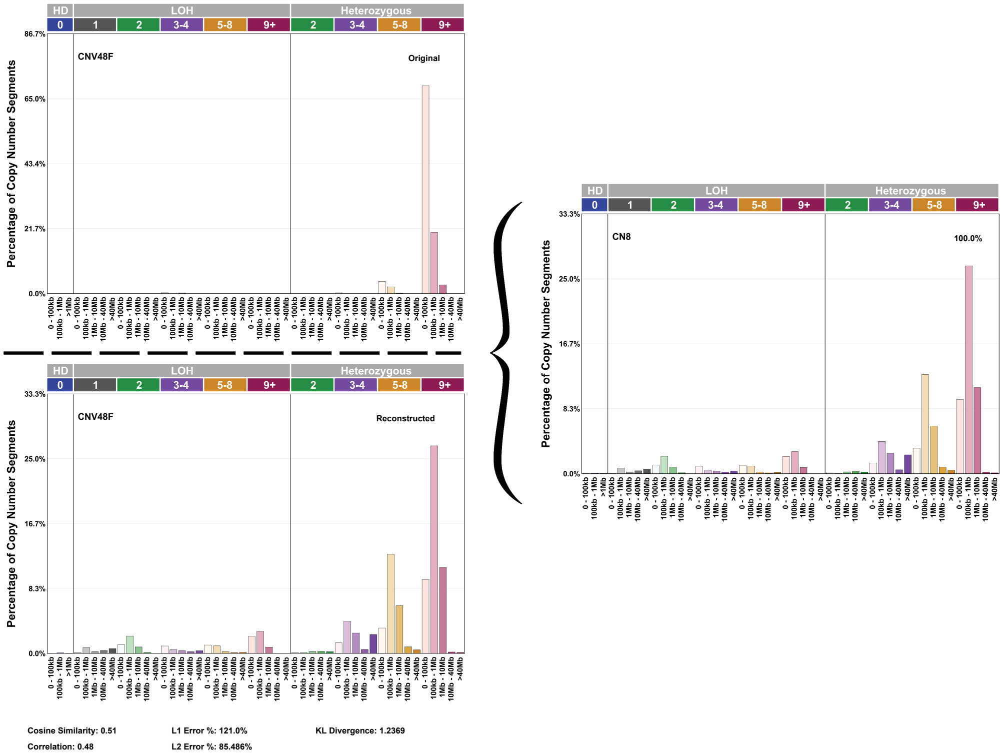
HD
LOH
Heterozygous
CNV48F
Original
Reconstructed
CN8
100.0%
Percentage of Copy Number Segments
Cosine Similarity: 0.51
Correlation: 0.48
L1 Error %: 121.0%
L2 Error %: 85.486%
KL Divergence: 1.2369

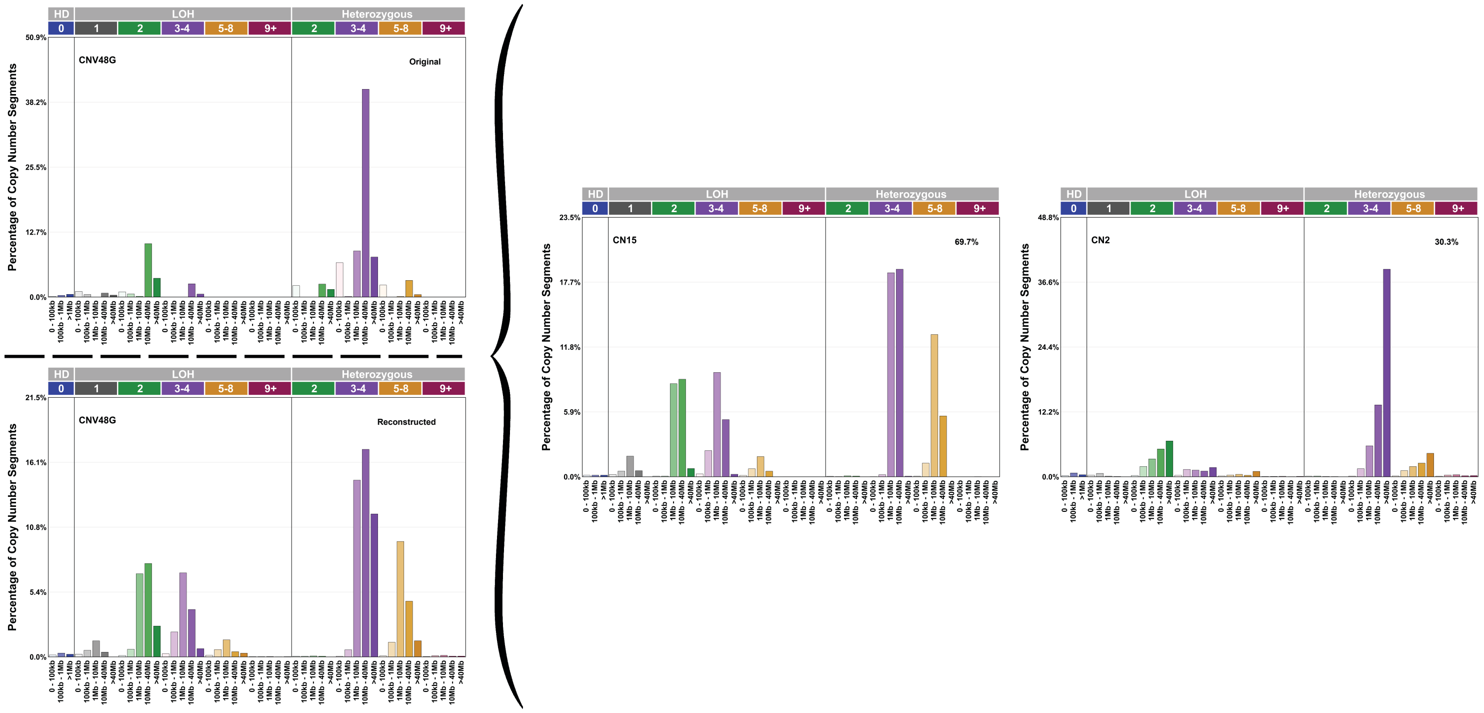


Cosine Similarity: 0.755 L1 Error %: 91.0% KL Divergence: 1.0878

Correlation: 0.723 L2 Error %: 66.834%

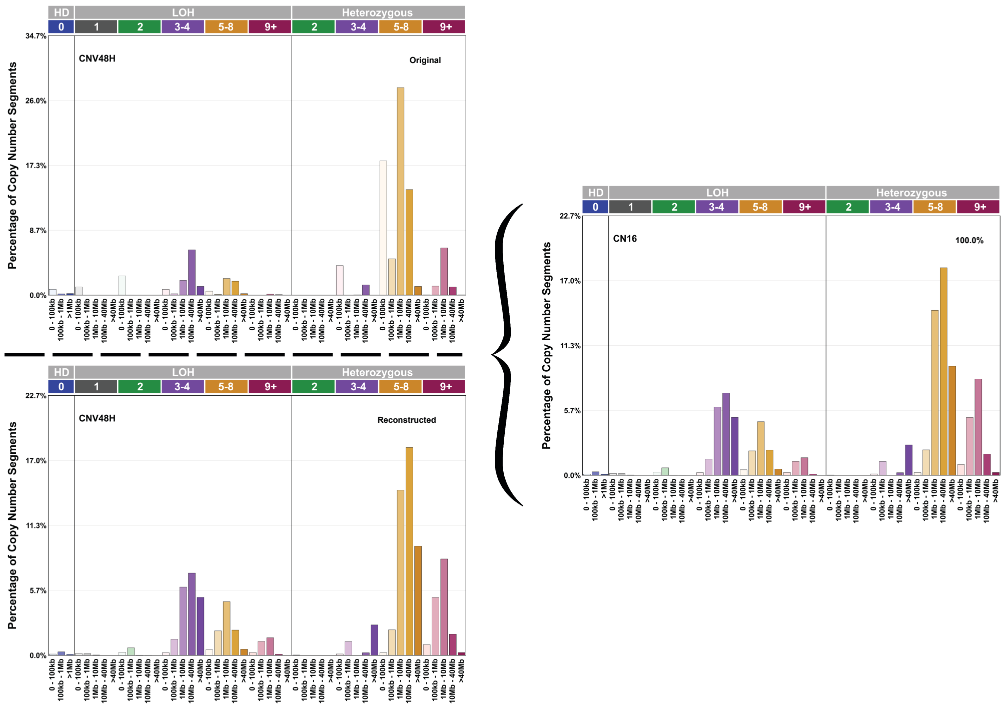


Cosine Similarity: 0.726 L1 Error %: 87.0% KL Divergence: 1000.0

Correlation: 0.669 L2 Error %: 68.644%

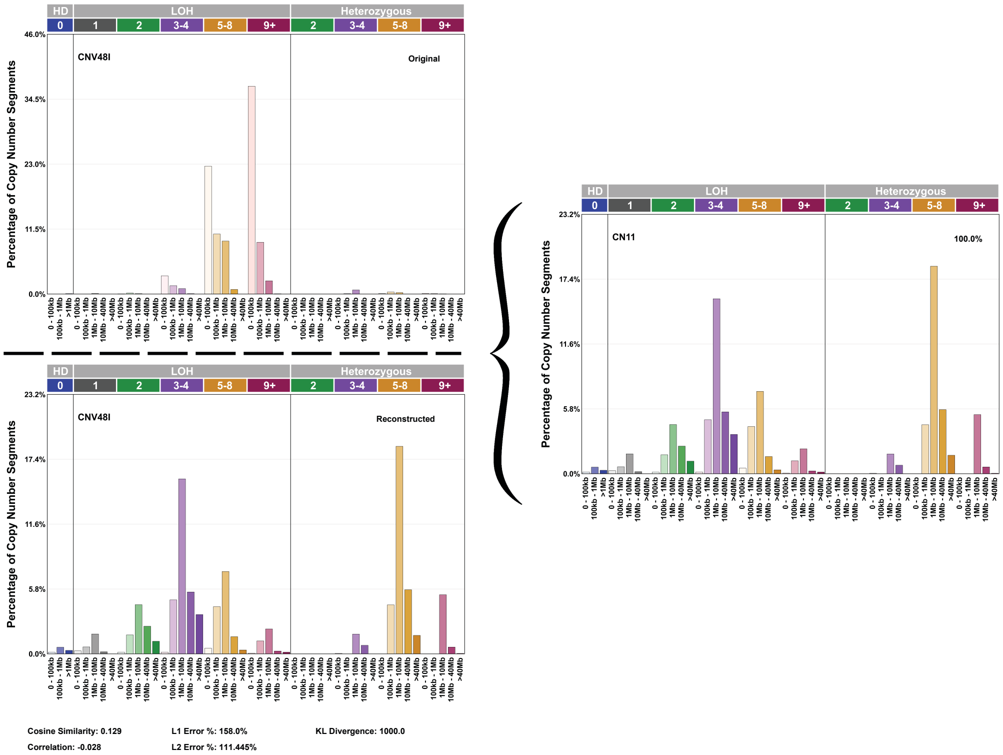
HD
LOH
Heterozygous
CNV48I
Original
Reconstructed
CN11
100.0%
Percentage of Copy Number Segments
Cosine Similarity: 0.129
Correlation: -0.028
L1 Error %: 158.0%
L2 Error %: 111.445%
KL Divergence: 1000.0

**c**

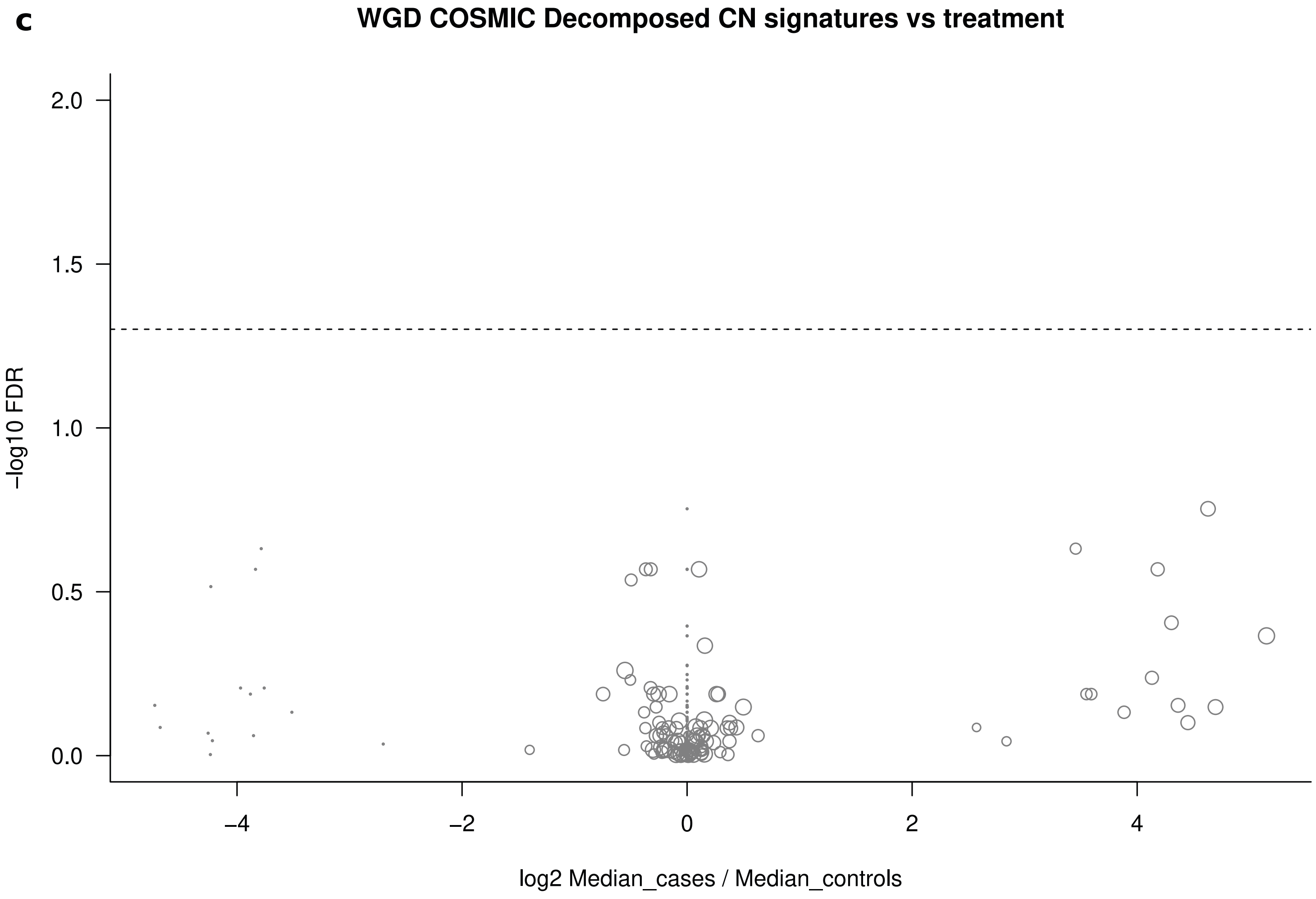
WGD COSMIC Decomposed CN signatures vs treatment
−log10 FDR
log2 Median_cases / Median_controls

**Figure S3. Comparison with CN signatures**

a) Comparison of the activity of CN signatures extracted de novo from HMF WGD tumors between samples exposed and unexposed to different anticancer treatments. No signature appears with significantly different activity between exposed and unexposed WGD tumors.

b) Equivalence (linear combination reconstruction) between CN signatures extracted de novo from the HMF cohort and CN signatures previously extracted from primary tumors (ref. 21).

c) Comparison of the activity of CN signatures extracted from primary tumors (ref. 21) between samples exposed and unexposed to different anticancer treatments. No signature appears with significantly different activity between exposed and unexposed WGD tumors in the HMF cohort.

### Figure S4

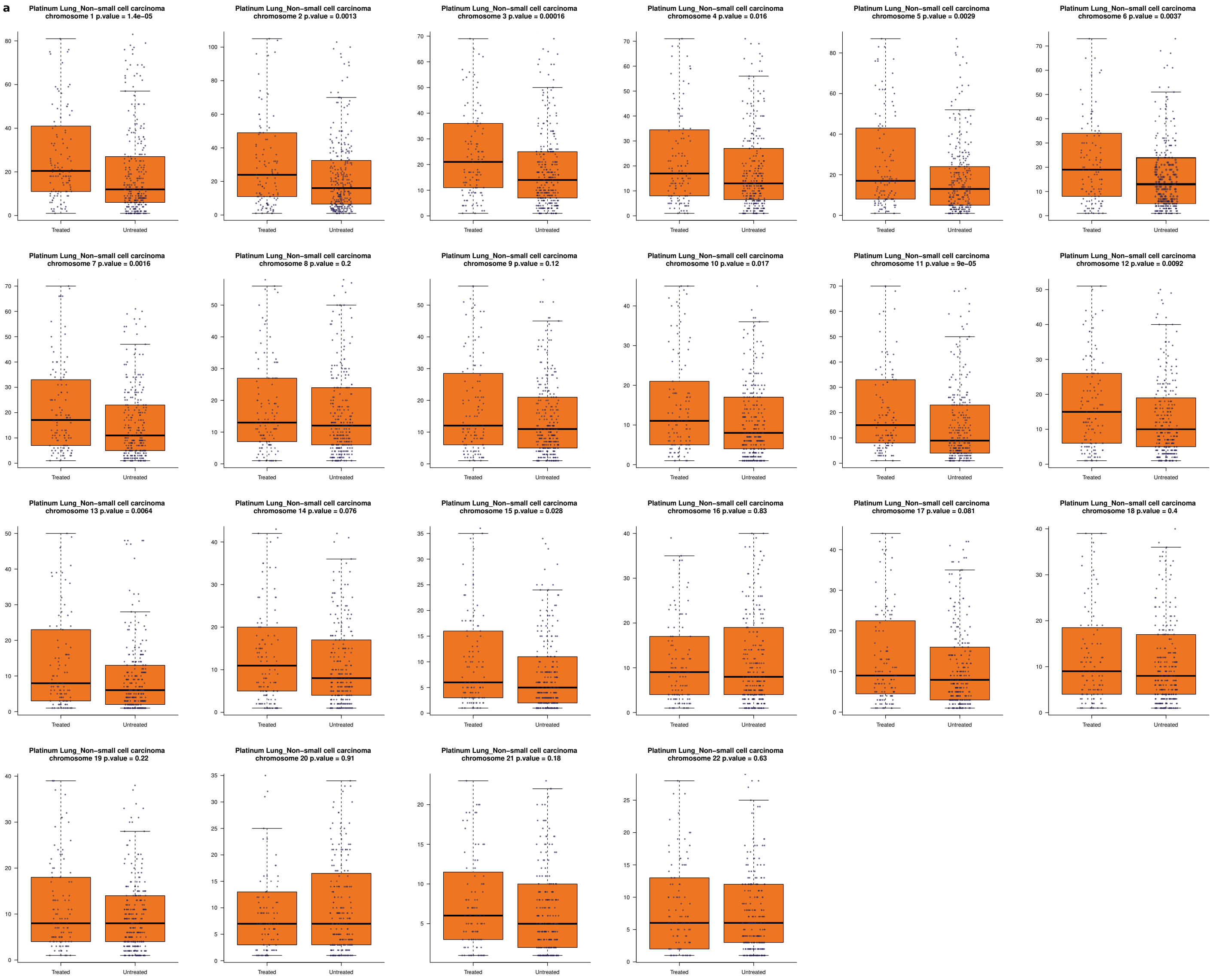
a
Platinum Lung_Non−small cell carcinoma chromosome 1 p.value = 1.4e−05
Platinum Lung_Non−small cell carcinoma chromosome 2 p.value = 0.0013
Platinum Lung_Non−small cell carcinoma chromosome 3 p.value = 0.00016
Platinum Lung_Non−small cell carcinoma chromosome 4 p.value = 0.016
Platinum Lung_Non−small cell carcinoma chromosome 5 p.value = 0.0029
Platinum Lung_Non−small cell carcinoma chromosome 6 p.value = 0.0037
Platinum Lung_Non−small cell carcinoma chromosome 7 p.value = 0.0016
Platinum Lung_Non−small cell carcinoma chromosome 8 p.value = 0.2
Platinum Lung_Non−small cell carcinoma chromosome 9 p.value = 0.12
Platinum Lung_Non−small cell carcinoma chromosome 10 p.value = 0.017
Platinum Lung_Non−small cell carcinoma chromosome 11 p.value = 9e−05
Platinum Lung_Non−small cell carcinoma chromosome 12 p.value = 0.0092
Platinum Lung_Non−small cell carcinoma chromosome 13 p.value = 0.0064
Platinum Lung_Non−small cell carcinoma chromosome 14 p.value = 0.076
Platinum Lung_Non−small cell carcinoma chromosome 15 p.value = 0.028
Platinum Lung_Non−small cell carcinoma chromosome 16 p.value = 0.83
Platinum Lung_Non−small cell carcinoma chromosome 17 p.value = 0.081
Platinum Lung_Non−small cell carcinoma chromosome 18 p.value = 0.4
Platinum Lung_Non−small cell carcinoma chromosome 19 p.value = 0.22
Platinum Lung_Non−small cell carcinoma chromosome 20 p.value = 0.91
Platinum Lung_Non−small cell carcinoma chromosome 21 p.value = 0.18
Platinum Lung_Non−small cell carcinoma chromosome 22 p.value = 0.63
Treated
Untreated

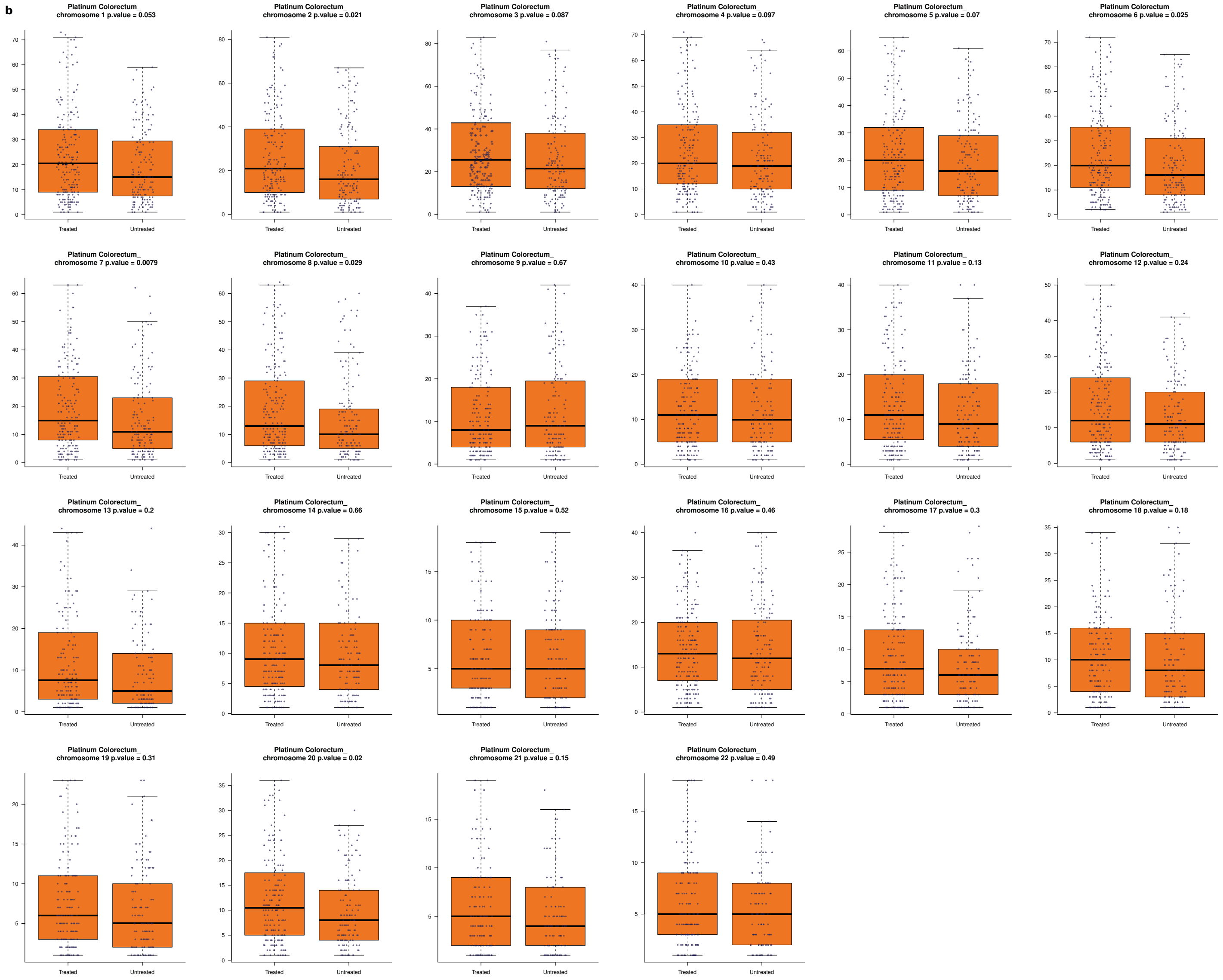


**Figure S4. Platinum CN footprint distribution across chromosomes.** Number of chromosomal fragments with copy number 1-4 and length smaller than 10 Mb across WGD lung (a) and colorectal (b) tumors exposed or unexposed to platinum-based drugs. Individual points represent the number of chromosomal fragments in a chromosome in an exposed or unexposed tumor. P-values represent the significance of a two-tailed Wilcoxon-Mann-Whitney test.

#### Figure S5

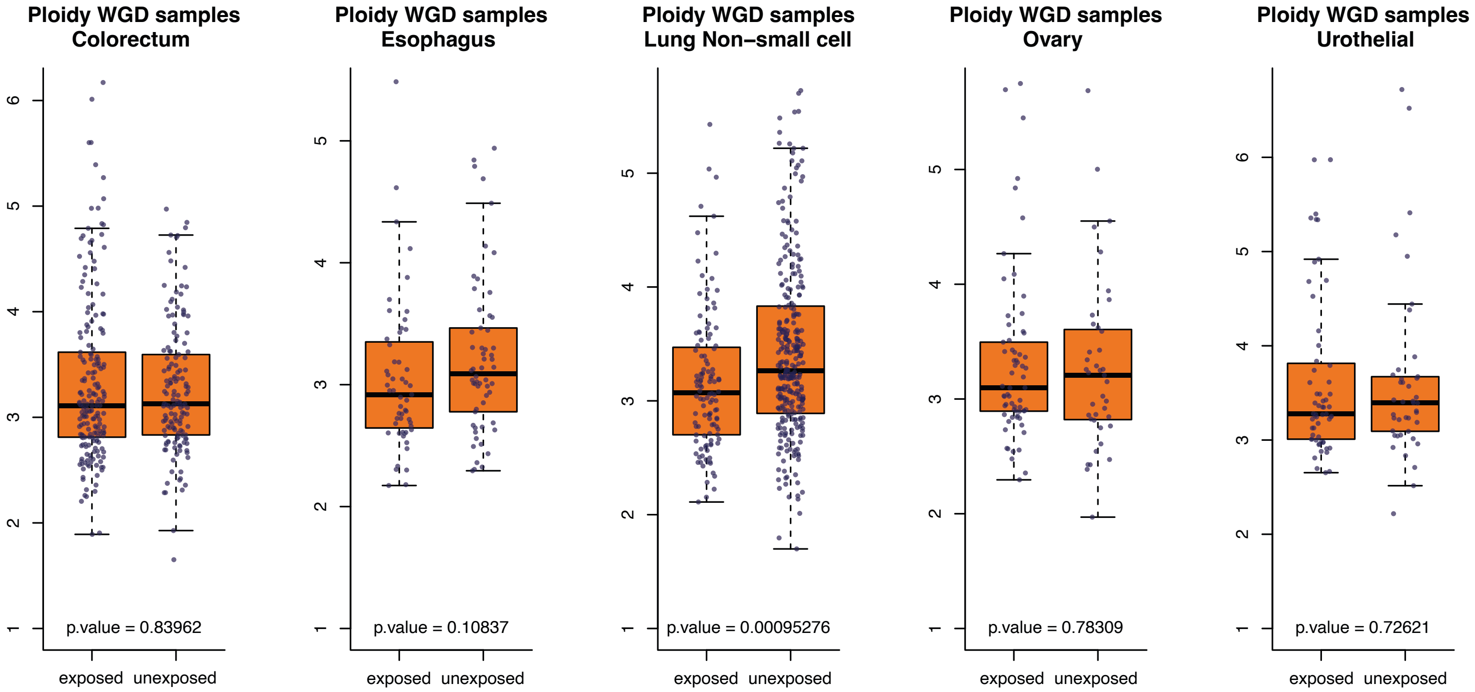


**Figure S5. Distribution of average ploidy of WGD tumors from different organs of origin that were exposed or unexposed to platinum-based therapies.** The distributions of both groups of tumors from each organ are compared using a two-tailed Wilcoxon-Mann-Whitney test.

Figure S6

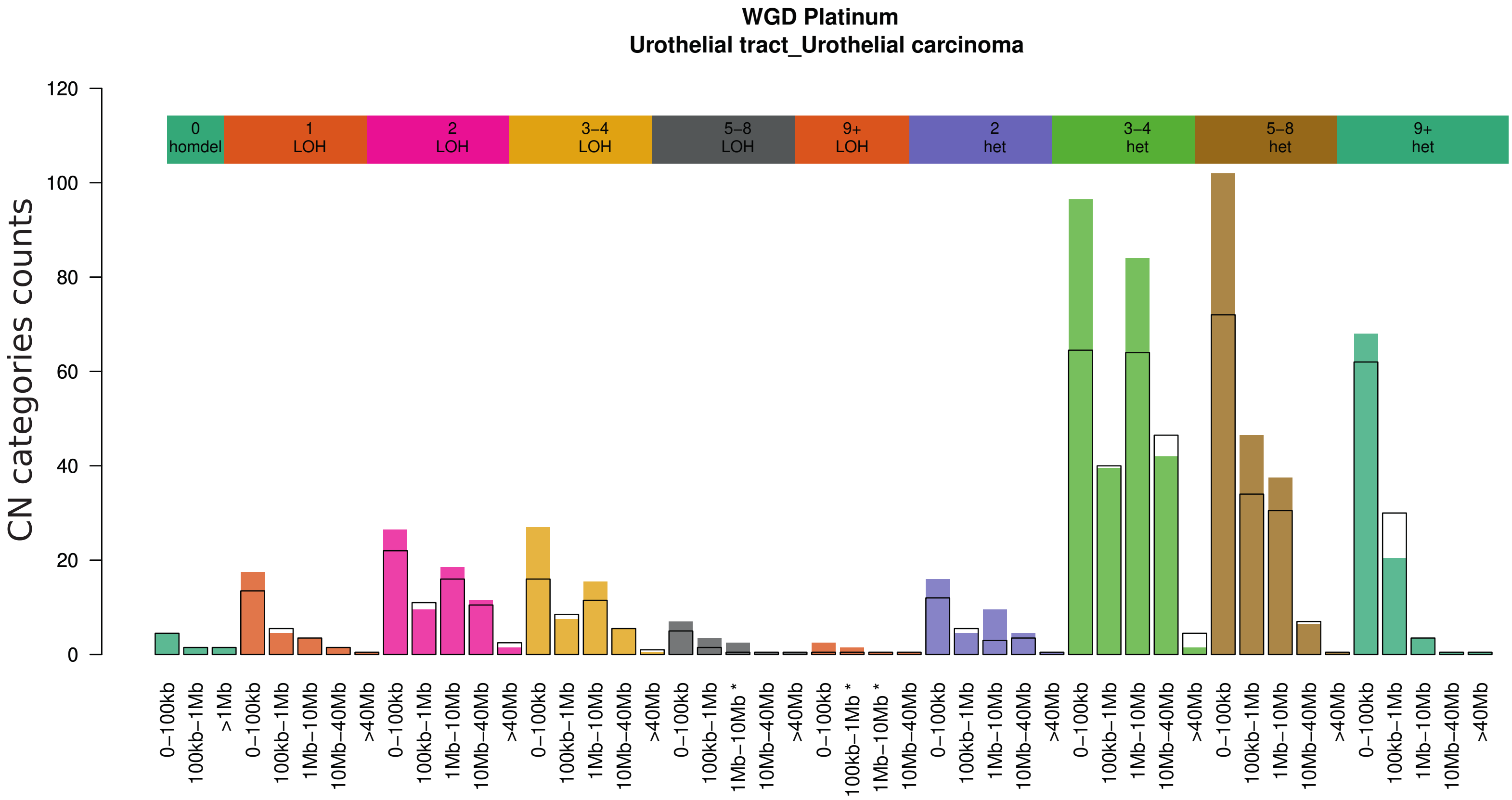
WGD Platinum
Urothelial tract_Urothelial carcinoma
CN categories counts
0 homdel
1 LOH
2 LOH
3–4 LOH
5–8 LOH
9+ LOH
2 het
3–4 het
5–8 het
9+ het
0–100kb
100kb–1Mb
>1Mb
100kb–1Mb *
1Mb–10Mb *
1Mb–10Mb
10Mb–40Mb
>40Mb

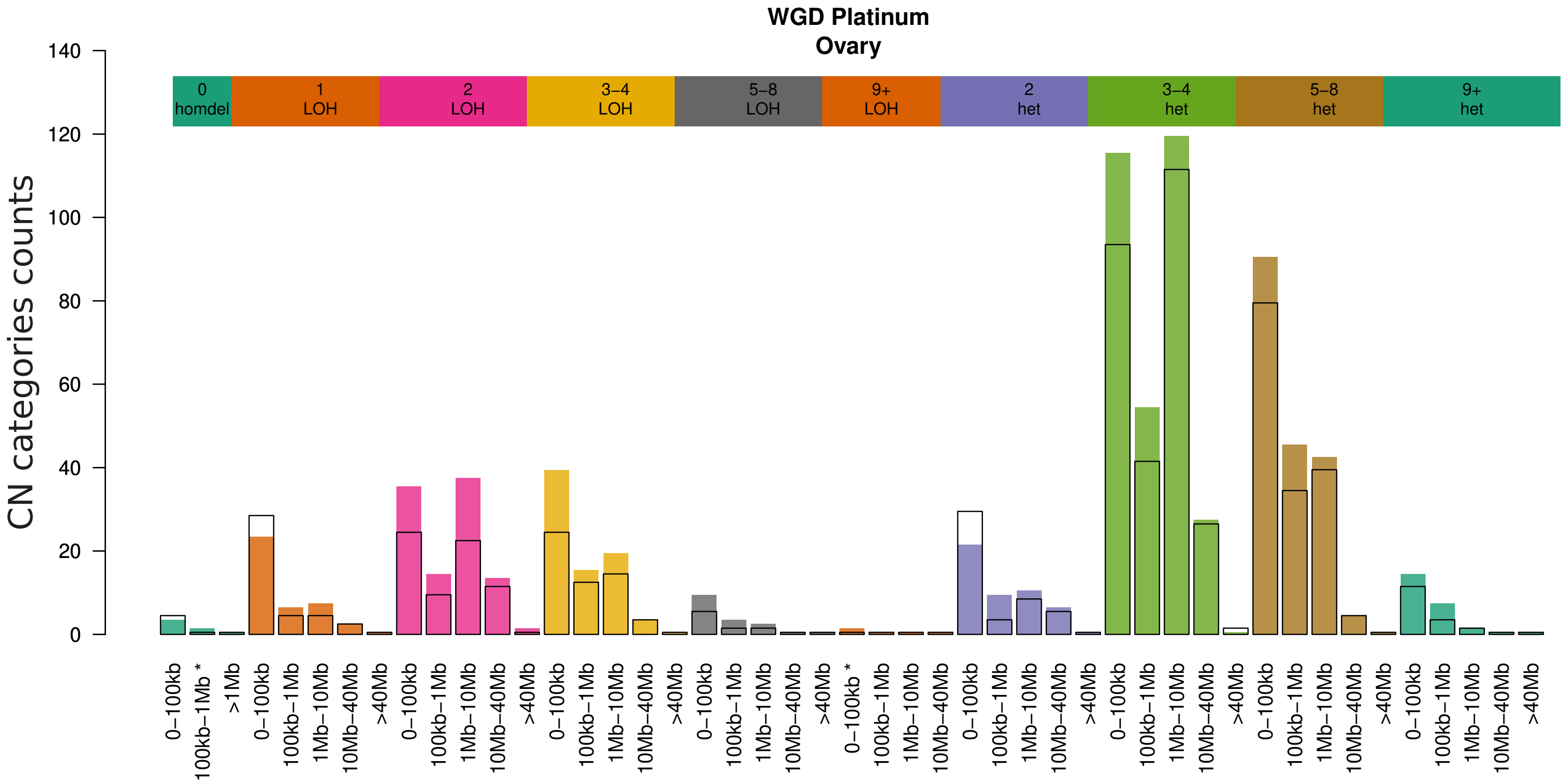
WGD Platinum
Ovary
CN categories counts
0 homdel
1 LOH
2 LOH
3–4 LOH
5–8 LOH
9+ LOH
2 het
3–4 het
5–8 het
9+ het
0–100kb
0–100kb *
100kb–1Mb
100kb–1Mb *
>1Mb
1Mb–10Mb
10Mb–40Mb
>40Mb

**Figure S6. Results of the comparison of CN categories across WGD urothelial and ovarian tumors exposed or unexposed to platinum-based therapies in the HMF cohort.**

#### Figure S7

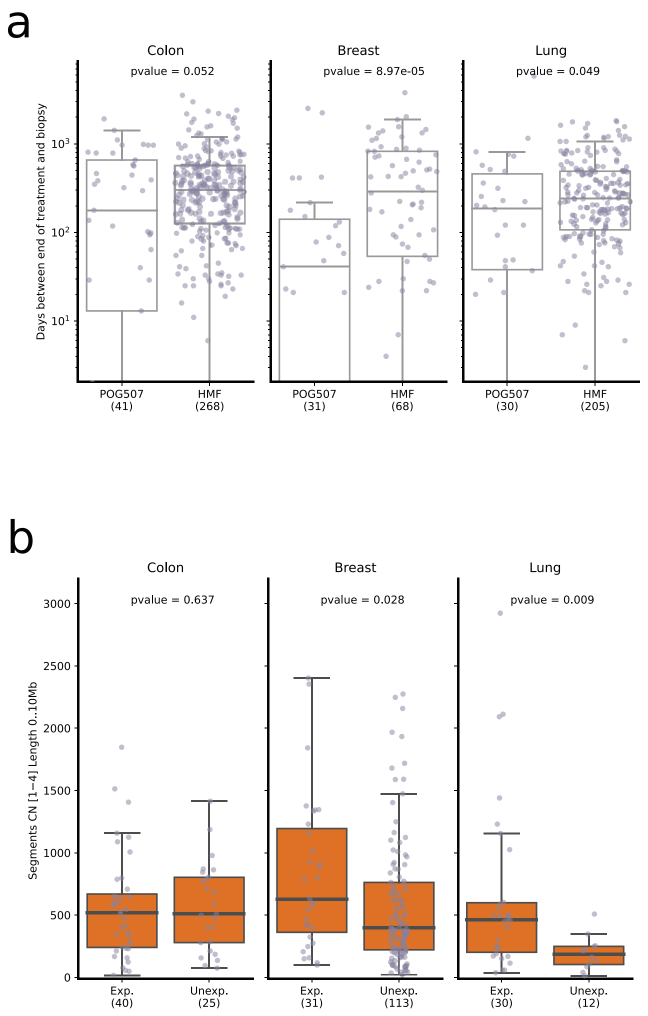


**Figure S7. Platinum CN footprint across tumors of an independent cohort (POG507).** a) Days elapsed between the end of platinum treatment and the biopsy of metastatic or recurrent tumors across the POG507 and HMF cohorts. Significantly longer time lapses for breast and lung tumors (and close to significant for colorectal tumors) are apparent across the HMF cohort. As a result, there is a higher likelihood of clonal expansion between treatment and biopsy (thus increasing the probability to detect platinum-related SBS and CN) in HMF metastatic tumors. This, together with the larger sample size probably explains why the platinum CN footprint is not as clear across POG507 colon tumors. One platinum exposed colorectal patient in the POG507 cohort did not have available CN data. b) Number of chromosomal fragments of size below 10 Mb identified across platinum-exposed and unexposed colon, breast and lung tumors. These numbers are significantly greater across exposed breast and lung tumors (two-tailed Wilcoxon-Mann-Whitney) and slightly greater (although not significant) across colon tumors. The platinum CN footprint is thus replicated in the POG507 cohort, except for colon tumors.

**Figure S8**

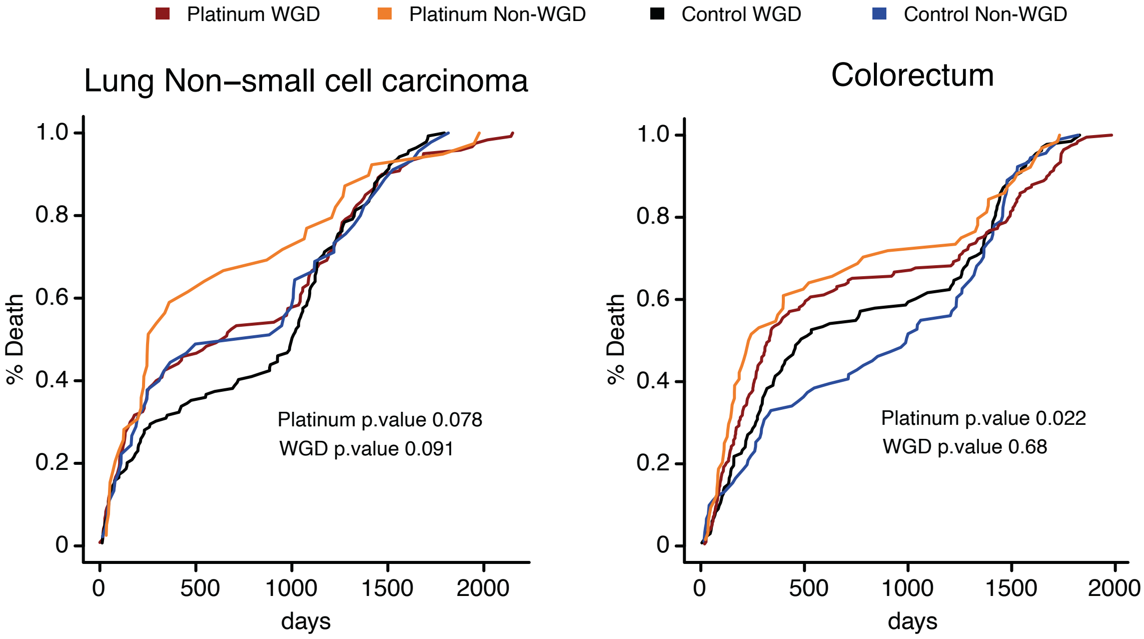


**Figure S8. Mortality curves of patients bearing metastatic tumors originated in the lung and colorectal.** Each curve represents a group of patients bearing either platinum-exposed or unexposed and WGD or non WGD tumors. The p-values were obtained using a Kaplan-Meier test.

### Supplementary Tables

**Table S1.** Fraction of WGD tumors across PCAWG and HMF cancer types.
**Table S2**. Number of WGD and non-WGD metastatic tumors of each tumor type across the HMF cohort exposed to different anti-cancer therapies.
**Table S3.** Association of CN features of WGD tumors with anticancer treatments across the HMF cohort.
**Table S4**. Overlap of tumors exposed to pairs of anticancer therapies in different cancer types across the HMF cohort.

### Supplementary Material of

#### Copy number footprints of platinum-based anticancer therapies

Santiago Gonzalez1, Nuria Lopez-Bigas1,2,3,^, Abel Gonzalez-Perez1,2,^

1. Institute for Research in Biomedicine (IRB Barcelona), The Barcelona Institute of Science and Technology, Baldiri Reixac, 10, 08028 Barcelona, Spain.
2. Research Program on Biomedical Informatics, Universitat Pompeu Fabra, Barcelona, Catalonia, Spain.
3. Institució Catalana de Recerca i Estudis Avançats (ICREA), Barcelona, Spain

^Corresponding authors
Abel Gonzalez-Perez:
Nuria Lopez-Bigas:
