## Supplement Response Reviewers for "Copy number footprints of platinum-based anticancer therapies"

### Full Revision

Review COMMONS

**Manuscript number:** RC-2022-01524
**Corresponding author(s):** Abel Gonzalez-Perez

#### 1. General Statements

We want to thank the three reviewers for their thorough revisions provided, which have helped us improve the manuscript. Following their comments, we have pursued two new main lines of analysis, the results of which are included in the new version of the manuscript.

- We have now validated the signal of platinum CN footprints observed across tumor types in the Hartwig Medical Foundation cohort in an independent cohort of metastatic tumors (POG570). Despite differences between these two cohorts, the same signal of increased chromosomal fragments of size below 10 Mb is observed in two tumor types.
- We have extended the analysis of CN signatures to those recently identified by Steele et al., 2022 (https://www.nature.com/articles/s41586-022-04738-6) across primary tumors. This new analysis revealed that no CN signature shows significant different activity between tumors exposed or unexposed to most frequent anticancer therapies. A simple measurement –i.e., the number of CN fragments of a range of lengths– proves in this case more effective than more complex CN signatures that capture combinations of CN features to identify the signal of exposure to platinum.

Other minor points raised by the three reviewers have also been addressed in the new version of the manuscript (see below).

#### 2. Point-by-point description of the revisions

Reviewer #1 (Evidence, reproducibility and clarity (Required)):

In this study, the author examine WGS data from two cohorts of cancer samples from previous studies: PCAWG, mostly representing primary tumors, and HMF, representing metastatic tumors. The HMF dataset represented 4902 metastatic tumors (including 2709 whole-genome doubling, i.e., WGD, metastatic tumors) from patients who had been exposed to 85 anticancer therapies. The study identified a pattern of large LOH events associated with the exposure of tumors of some cancer types to platinum-based therapies. This pattern is characterized by a significant increase in the number of chromosomal fragments in exposed tumors with respect to their unexposed counterparts. The findings could support the hypothesis that WGD may provide tumors with an advantage to withstand the effects of structural variation.

We thank the reviewer for their accurate summarization of our manuscript.

Specific comments:

1. There are a number of statements that would suggest that there is some uncertainty regarding the robustness of results, and that the analysis of additional cohorts may be needed to substantiate the overall findings. For example, page 4: "It is also plausible that more numerous cohorts of exposed tumors are required to understand whether the observed differences are indeed robust." Page 5: "Further analysis with larger cohorts are required to clarify this point, which appears especially to clarify whether a significant imbalance in favor of deleted chromosomal fragments does occur across platinum-exposed lung tumors." However, the abstract does not seem to reflect this level of uncertainty in reporting the main findings.

We thank the reviewer for pointing this out. It is important to highlight that the first statement cited by the reviewer refers to the potential taxanes-related CN pattern, which is not mentioned in the abstract. The second statement refers to the fact that while we observe no significant differences of ploidy between platinum exposed and not exposed WGD tumors, we caution that this may change in larger cohorts. Following the reviewer's observation, we have now deleted this sentence from the abstract. Therefore, in the current version of the manuscript, all statements presented in the abstract are robustly observed across tumors.

Moreover, we have now replicated the finding of the platinum CN footprint across the POG570 cohort (https://www.nature.com/articles/s43018-020-0050-6), the second largest cohort of whole-genome sequenced metastatic tumors available. (See answers below for details.)

2. Many of the findings made appear to apply not to all tumors but are found within tumors of specific cancer types. However, the abstract does not appear to note this.

As stated in the previous point, all statements contained in the abstract in the current version of the manuscript are replicated across platinum-exposed tumors of different cancer types across the Hartwig Medical Foundation metastatic cohort. Results on different tumor types with WGD samples exposed to platinum treatment are presented in Figures 2d and 3c and Supplementary Figure 5. A summary of the platinum CN footprint resulting from the aggregation of platinum CN patterns observed across all WGD exposed tumors is presented in Figure 3a.

We consider precisely this robustness of the CN pattern observed across platinum-exposed tumors of different cancer types as evidence supporting the proposed universal platinum CN footprint. Conversely, we do not propose a taxane-related CN footprint, precisely due to the lack of such robustness, as explained in the manuscript. We haver rewritten this part of the manuscript to make this point clearer.

Finally, as mentioned above, we have now replicated the platinum CN footprint across platinum-exposed tumors in the POG570 cohort (https://www.nature.com/articles/s43018-020-0050-6). Despite the limitations in sample size and the overall shorter time elapsed between the end of the treatment and the biopsy of the metastasis compared to the HMF cohort, as well as a different method for CN calling 2 out of 3 tumor types (breast and lung) with at least 10 platinum exposed and unexposed samples show exactly the same footprint while comparing platinum treated vs untreated. (See details below.)

3. With regards to additional cohorts, there is a POG570 cohort of WGS data on 570 recurrent or metastatic tumors (Nature Cancer 2020, PMID: 35121966), some 82% of which were from patients receiving systemic therapy before biopsy. Is it possible that some of the patterns identified using the HMF datasets could be validated in the POG570 datasets? If not, what numbers of tumors would be needed for the patterns of interest to be reliably identified?

We thank the reviewer for pointing us in the direction of this very interesting dataset. We have now analyzed the POG570 cohort (https://www.nature.com/articles/s43018-020-0050-6) using the clinical and treatment data and chromosomal fragment calls provided by the authors. Briefly, for each tumor, we computed the length of each chromosomal fragment identified and then counted the number of fragments with copy number 1-4 and length below 10 Mb, which constitutes our measurement of the intensity of the platinum CN footprint. Then, we compared the number of these fragments across platinum exposed and unexposed colon, breast and lung tumors. We selected these tumor types because only for them are there at least 10 exposed and unexposed tumors in the POG570 cohort.

The results of this comparison are shown in the new Figure S7a of the manuscript, which we reproduce here.

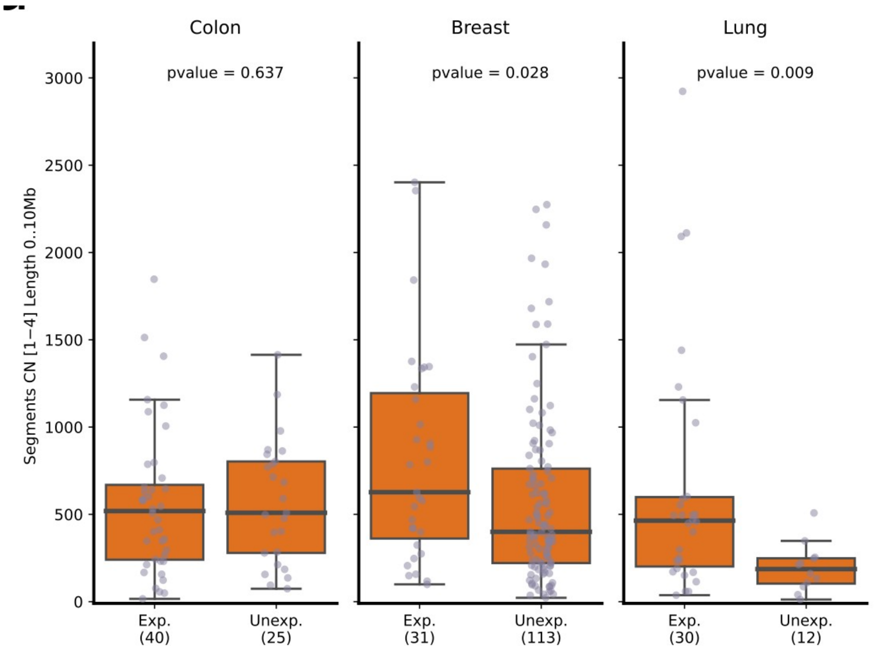


We found that platinum-exposed breast and lung tumors have a significantly higher number of chromosomal fragments of CN 1-4 and length below 10Mb than their unexposed counterparts, thus replicating our observation across HMF tumors. In the case of colon tumors the difference is not significant. Several differences in the composition of the cohorts and the analysis of the data must be taken into account in the analysis of these results. On the side of the data, the calls of chromosomal fragments have been done using different algorithms. On the side of cohorts composition, one important difference is the number of days elapsed between the end of the exposure of the patients to platinum and the moment of the biopsy of the metastatic or recurrent tumor. The differences between the three tumor types analyzed across both cohorts are now represented in Figure S7b of the manuscript, reproduced here.

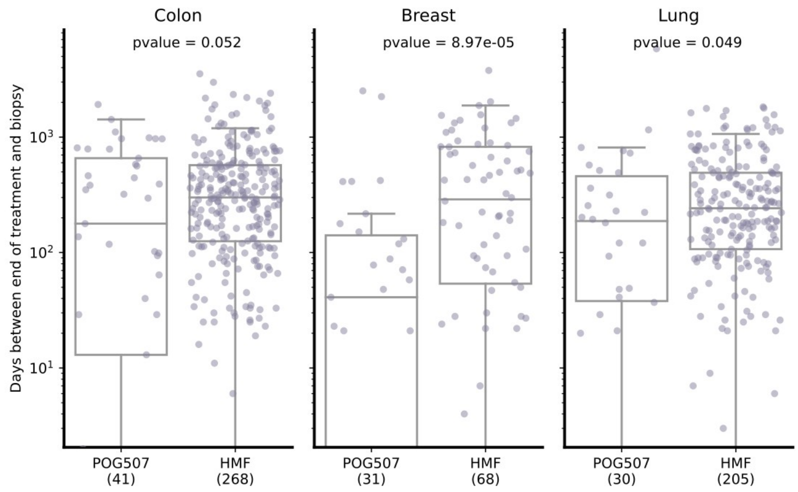


Across the HMF cohort, colorectal tumor patients exposed to platinum have a median of 300 days between end of treatment and biopsy of the metastasis, but only 178 across POG570 colon tumors (difference close to significance). The same gap is appreciable across breast tumors (289 days vs 41 days; significantly higher across HMF patients) and lung tumors (242 days vs 187.5 days; borderline significantly higher across HMF patients).

In a previous work (https://www.nature.com/articles/s41467-021-24858-3) we demonstrated that longer time elapsed between the end of the treatment and the biopsy accounted for a higher likelihood of a full clonal expansion upon treatment and, as a consequence a higher probability to detect the SBS platinum footprint through bulk sequencing. Since the same must apply to the CN footprint, the significantly shorter time between the end of treatment and biopsy across POG570 tumors would make its detection more difficult. Nevertheless, it is still nicely reproduced at least across breast and lung tumors.

In summary, despite the differences between the HMF and POG570 cohorts, the platinum CN footprint is replicated across tumors of the latter, thus providing independent support to its presence in platinum-exposed tumors.

4. The PCAWG cohort is described as comprising all primary tumors, but in fact there are some metastatic tumors in PCAWG cohort. In particular, most of the TCGA melanoma (SKCM) samples are metastatic (PMID: 30401717). This may have bearing on using comparisons between PCAWG and HMF as a surrogate for primary versus metastases.

The reviewer is correct that there is a very minor representation of metastatic tumors in the PCAWG cohort, specifically, across melanomas. It is nevertheless a good choice for a whole-genome sequenced cohort of tumors, which has been used repeatedly in primary-to-metastatic comparisons (see, for example, PMID: 31645765; https://www.biorxiv.org/content/10.1101/2022.06.17.496528v1). Since the vast majority of tumors in the PCAWG cohort are primary, and the comparisons presented in Figure 1 encompass the entire cohort (with the exception of Fig. 1b, where melanomas are not included), the influence of a few tumors are not expected to confound the results. In any case, differences between primary and metastatic tumors are still very apparent in Figure 1c; exclusion of PCAWG melanomas, if anything, would make them still more apparent.

5. For each boxplot, the number of tumors represented in each group should be indicated somewhere (e.g., along the bottom).

We thank the reviewer for bringing this oversight to our attention. We have now added the number of tumors in each group in all relevant figures. Specifically, the number of tumors in each group appear now in Figures 1c and d, 2d and e, 3 c and d, and 4 a and b, as well as in relevant Supplementary Figures.

6. For Figure 1a, is a color legend needed here?

Thanks for bringing this to our attention. We have now added a color legend to represent WGD and near diploid (non-WGD) tumors in this panel.

7. For analyses comparing HMF to PCAWG (e.g., Figure 1c), the p-values ought to corrected for cancer type (e.g., using a linear regression model with cancer type as a factor).

We thank the reviewer for the suggestion to look at the influence of tumor types in the observed differences in the fraction of LoH tumors in primary and metastatic cohorts. Following their suggestion, we have now carried out separate comparisons of the overall ploidy and fraction of genome LoH of tumors per cancer type (Fig. S1b-i in the reviewed version of the manuscript). This analysis is, of course, limited to tumor types represented in both PCAWG and HMF cohorts. As in the pan-cancer analysis, no significant differences in ploidy are observed in any cancer type. Conversely, significant differences in LoH fraction appear in colorectal, prostate and kidney WGD primary and metastatic tumors. Regardless of statistical significance, in the majority of cancer types, the fraction of LoH genome across tumors appears greater in metastasis than in primaries. Therefore, our starting observation that more LoH is observed across metastatic than primary WGD tumors and that this could be related with the former’s exposure to anticancer therapies holds in this per-tumor-type analysis.

8. For Figure 1d, are the numbers of tumors in each category indicated in parentheses?

The reviewer is correct. We have clarified this in the caption of Figure 1d.

9. For figures 2d and 2e legend, the numbers of tumors in exposed vs unexposed groups for each category should be indicated. Similar for Figures 3a, 3c, 3d.

We thank the reviewer for pointing out this oversight. We have now added the numbers of exposed and unexposed tumors in the relevant plots in Figures 2 and 3. Moreover, we have added a new Supplementary Table (Table S2) with the numbers of tumors of each cancer type exposed and unexposed to each treatment across the HMF cohort.

10. For figure 2c, what is the statistical test used and multiple testing correction applied? Could this be noted in the figure legend?

Following the reviewer's observation, we have included the name of the test (Wilcoxon signed-rank test) in the caption of Figure 2c. The p-values shown are corrected for False Discovery Rate: this is now indicated in the Figure caption.

Reviewer #1 (Significance (Required)):

The study makes effective use of public genomic resources to make new observations regarding platinum-based anticancer therapies. The observations identify patterns within specific cancer types. The analysis is exploratory in nature and would benefit from independent observation in an independent cohort, though it is not clear whether such cohorts may exist in sufficient numbers.

As explained above in detail, motivated by this comment by the reviewer, we have now validated the platinum CN footprint across an independent cohort of metastatic tumors (POG570).

Reviewer #3 (Evidence, reproducibility and clarity (Required)):

Analyzing the genome wide copy number patterns across publicly available ~2700 primary and ~5000 metastatic tumors treated by a number of different classes of chemotherapy agents, the authors find a distinct signature of CNVs in tumors treated with platinum-based agents. These platinum-exposed tumors are characterized by a significant increase in the number of chromosomal fragments of lengths between 10 Kb-10 Mb, and this tendency correlates with dosage (approximated by previously published platinum induced mutational signatures). Also, it is interesting that comparison of WGD with non-WGD treated-vs-untreated samples shows that WGD samples tolerate larger CNVs, suggesting relaxed selection against large CNVs in WGD (or WGD as a mechanism to accumulate large CNVs).

Previous works have focused on mutational signatures of various environmental exposures and drugs. This paper attempts to extend the previous research by looking at patterns of copy number variations. The work is somewhat motivated (see comment below) and the experimental design and execution are reasonable.

The manuscript is well written. The method section could be elaborated more for reproducibility.

We thank the reviewer for their appreciative comments on our manuscript. Following their observation, we have carefully reviewed the methods section thinking on the reproducibility of our results.

Major comments:

1. Overall, the results are modest. Although statistically significant, the increase in specific classes of CNVs in treated v untreated WGD mets is modest (Fig 2d), casting a doubt on clinical significance.

While the statistical significance of the association of independent CN features to the exposure to platinum-based drugs is not as high as that observed for platinum-related single nucleotide variants (https://www.nature.com/articles/s41588-019-0525-5), this does not mean that the signal is modest. When the number of CN fragments of length below 10Mb with CN 1-4 of tumors exposed and unexposed to platinum (across three cancer types) are compared, the signal is very clear (Figure 4a). Across cancer types, these particular chromosomal fragments are more abundant (significantly in most cases) in platinum-exposed tumors than in their unexposed counterparts. The increase across exposed tumors is between 13% and 387%.

The number of CN fragments in this range of sizes that may be identified due to the exposure to platinum is much more stringently limited by the size of the genome than the corresponding number of SNVs. This is why while we observe thousands of platinum-related SNVs in exposed tumors(https://www.nature.com/articles/s41588-019-0525-5), the numbers of observed CN fragments are smaller, making the signal less strong. However, while each single nucleotide variant affects a single nucleotide, a chromosomal fragment in the middle of the range observed would affect thousands of base pairs. This makes the cumulative effect of the platinum CN footprint on exposed tumors and normal cells is much larger than that of single nucleotide variants. In other words, the effect of the exposure to platinum on the landscape of CN fragments, far from modest is more consequential than that of SNVs. Thus, while a clinical application of the identified platinum CN footprint is out of the scope of this work, we do believe that, like its mutational footprint counterpart (described in the abovementioned papers) it does have implications for chemotherapy survivors.

To clarify further the effect size of the exposure to platinum, following the reviewer's comment, we have now added a fold-change to the comparisons between exposed and unexposed tumors presented in figure 4a. Furthermore, we have added the following consideration to the last paragraph of the Discussion section:

*Moreover, these SVs –as described by the platinum CN footprint– are bound to affect much larger genomic portions than platinum-contributed point mutations (6, 11). Therefore, their impact on exposed healthy cells and on the development of late effects of the chemotherapy could potentially be greater than those caused by previously recognized platinum-related SBS footprints.*

2. Three of the four drugs that yielded significant patterns seems to have largest sample sizes (Fig 2a). Is there a link between sample size and detection power? In general, robustness of the signals is not analyzed, relative to subsampling of tumors or genomic regions etc. Indeed, the authors have noted the potential lack of robustness somewhere in the manuscript.

The reviewer is right that the sample size is an important limiting factor for the detection of CN patterns related to anticancer therapies. This is much more acute than in the case of footprints of single base substitutions, thousands of which are contributed by platinum (for example) to the genome of exposed cells. In contrast, limited by their sheer size, only a few dozen extra chromosomal fragments are contributed by the same treatment to metastatic tumors. This is the reason why, rather than carrying out a subsampling exercise, we have resorted to identifying CN patterns in the tumors of different tumor types separately. As suggested by the reviewer, we have only considered robust the CN pattern that is detected across all cancer types with exposed samples, that is, that of platinum-based therapies.

Adding robustness to the detected platinum CN footprint, we have now replicated its finding in a totally independent cohort of tumors, the POG570 cohort. See above answer to point 3 raised by reviewer 1.

Moreover, we include a paragraph in the Discussion section dedicated to comment on the question of power for the detection of the CN footprints associated to other therapies.

3. Given inter-individual heterogeneity, analyzing longitudinal data of pre-treated primary and post-treated mets rom same individuals would really help strengthen the findings.

We agree with the reviewer that such a comparison would be very interesting. Unfortunately, pretreatment samples of the primary tumors of patients in the Hartwig Medical Foundation cohort are not available.

4. The authors show that several CN features show and increase upon platinum treatment. Are these independent observations? A global correlation among the 48 features in various samples classes (WGD/non-WGD, Primary/Met, treated/untreated) should be done and equivalence class of features defined. Otherwise, the biological significance of these observations could be overstated.

The reviewer is right that there is correlation between the CN features used to identify the CN footprint. Nevertheless, these features, which have been defined elsewhere (https://www.nature.com/articles/s41586-022-04738-6) for the identification of CN signatures are only used in the context of our analysis to determine that some of them (despite their potential correlation) are different between platinum exposed and unexposed tumors. Precisely, taking into account the correlations between CN features, our conceptualization of the platinum CN footprint is the number of chromosomal fragments with copy number between 1 and 4 with length below 10 Mb. Moreover, using this definition and not the original CN features, we are able to replicate the observation of the platinum CN footprint across an independent cohort, which provides further robustness to its identification.

In our work, in summary, the CN features are only a means to the end of identifying a quantitative difference in the structure of chromosomal fragments between tumors exposed or unexposed to a certain anticancer therapy.

5. The only mechanistic link between platinum treatment and the observed CNV patterns is speculated to be via platinum-induced DNA breaks and errors during correction. This seems like a very general mechanisms applicable to any exposures (environmental or drug) that induces breaks. This lack of specificity makes it hard for me to understand the rationale to study CNV patterns - why, after all, should one expect to see a CNV signature?

The question posed by the reviewer –are there any treatment related CN footprints?– is precisely the starting point of our study. We thus carried out an unbiased discovery of CN patterns related with the exposure to different treatments. Upon identification of the association between the exposure to platinum and the increase of LoH chromosomal fragments of 10kb-10Mb signatures with different copy number, we hypothesize that the platinum-induced increase of double strand breaks and their faulty repair may be the underlying mechanism. We absolutely agree with the reviewer that other therapies inducing double strand breaks could lead to a similar –or other– CN footprint. Nevertheless, we have not been able to detect other consistent CN footprint associated with any anticancer therapy across tumors in the Hartwig Medical Foundation cohort. Whether this is due to the lack of statistical power or some underlying mechanistic difference between platinum-based and other drugs (see for example the causes underlying differences in the detectability of platinum and 5FU-related footprints; https://www.nature.com/articles/s41467-021-24858-3) we are not currently able to answer.

6. Authors should contrast their findings with those in https://www.nature.com/articles/s41586-022-04738-6.pdf

We thank the reviewer for this suggestion. Actually, taking advantage of the fact that the tool employed to extract CN signatures de novo (the original SigProfiler aimed at mutational signatures extended to CN features) was available prior to the publication of this article, we had already carried out a CN signatures extraction de novo from the HMF cohort. We then asked if any of these CN signatures (their activity across tumors) is significantly associated with the treatments in the cohort, and found none (current Fig. S3a). In the manuscript we hypothesize that this is due to the intrinsic difficulties in defining CN signatures, as opposed to SBS and DBS signatures. This is why we decided in our study to focus on a collection of individual CN features that show differences between platinum-exposed and unexposed tumors to define the platinum CN footprint.

Following the suggestion by this (and other) reviewer, we have now carried out the same analysis (identification of CN signatures potentially related to exposure to anticancer therapies) using the set of CN signatures originally defined by Steele et al in their paper (reference 21 in our manuscript). This analysis yields negative results too (current Fig. S3c). We also now include the equivalence (established through linear reconstruction) between the CN signatures extracted de novo from the HMF cohort and the CN signatures originally defined by Steele et al. across primary tumors. (This equivalence is provided by the SigProfiler upon extraction.) In general, the signatures extracted across HMF tumors bear little resemblance to those extracted from primary tumors (highest cosine similarity of a linearly reconstructed signature, 0.775). This is presented in current Figure S3b.

Taken together, these results further strengthen our point that a CN footprint defined from differences in individual CN features are probably more appropriate than CN signatures in their current format to identify the effects of anticancer therapies on the CN landscape of exposed cells.

7. For pyrimidine treated samples... "significance is lost across non-overlapping tumors". Authors should ascertain that this is not simply a matter of power. Also, would the significance not be lost for non-overlapping platinum-treated samples?

We thank the reviewer for pointing out the lack of clarity in our statement. To solve it, we have included a new Supplementary Table (Table S4) containing the number of WGD tumors exposed to different pairs of anticancer therapies across cancer types in the HMF cohort.

Let’s look at three cancer types showing the platinum CN footprint with different degrees of overlap of platinum and pyrimidine analogs exposed tumors. In the case of colorectal tumors,

194 out of 220 pyrimidine analogs exposed WGD tumors are also exposed to platinum. No signal is observed when the 26 tumors solely exposed to pyrimidine analogs are compared to tumors that are unexposed to pyrimidine or platinum (as shown in the Figure below, the p-values of which correspond to a two-tailed Wilcoxon-Mann-Whitney test).

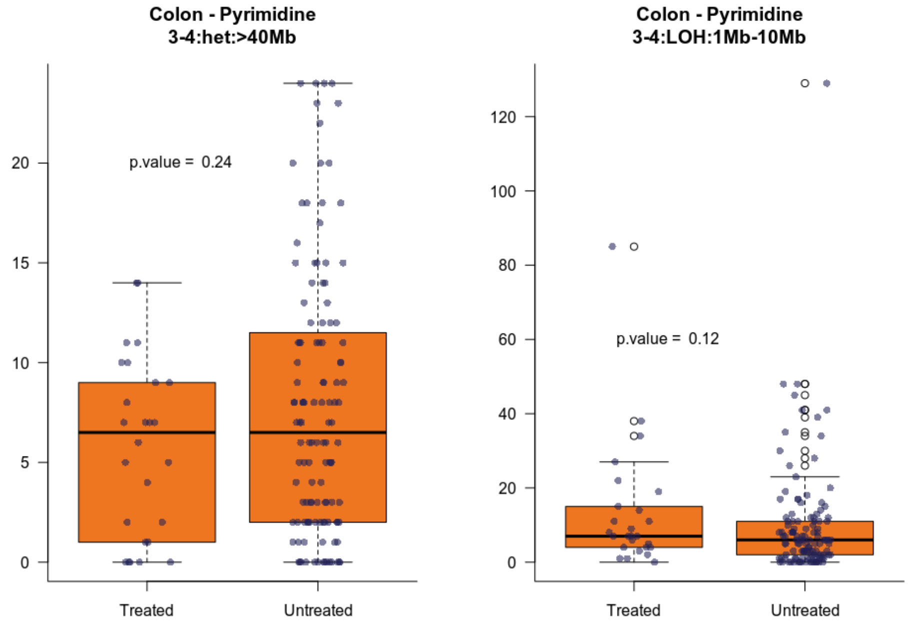


The reverse analysis is impossible, as only 4 WGD tumors are exposed to platinum but not to pyrimidine analogs. In the case of lung tumors, the 22 tumors exposed to platinum and pyrimidine analogs constitute the entire pyrimidine analogs exposed set. When only the 84 tumors exposed solely to platinum are compared to tumors unexposed to platinum or pyrimidine analogs, CN features associated to platinum exposure still appear different. Finally, in the case of ovarian tumors, no exposure to pyrimidine analogs is recorded, as it is not employed in the treatment of this malignancy.

In summary, the significance of platinum-related CN features is present when only platinum-exposed tumors are included in the comparison. The signal observed is thus attributable to the exposure to platinum-based drugs. The number of exclusively pyrimidine-exposed tumors are few across tumor types, and thus at this stage we are not able to rule out the existence of a pyrimidine associated footprint.

8. Interpretation of Fig 3b. "Had this increase in the number of 10Kb-10Mb chromosomal fragments across exposed tumors arisen through positive selection, we would expect to observe a concentration at specific genomic regions containing resistance genes." This needs to be tested specifically. A "concentration" perhaps would not jump out in a visual inspection of the global pattern, which does seem to show variability.

Following the reviewer's suggestion, we have now compared the number of chromosomal fragments of CN 1-4 and size smaller than 10 Mb observed in each chromosome across platinum exposed or unexposed lung and colorectal tumors. The results of these comparisons are presented in Figure S4a,b. This figure shows that more fragments of this size range are observed for all chromosomes across exposed tumors than across their unexposed counterparts. In most cases the differences are significant. This means that the increase of chromosomal fragments of size below 10Mb is not restricted to one or few chromosomes. It is rather a general effect distributed along the entire genome.

Minor comments:

9. "the ploidy of tumors with WGD varies in a range between 2.9 and 3.6 (quartiles 1 and 3, Fig. 1a)". I am not sure, there seem to several orange (WGD) points with ploidy below 2.9.

The reviewer is correct that several WGD tumors possess a ploidy below 2.9. This is because the cited values 2.9 and 3.6 correspond, respectively to the lower and upper limit of the first and third quartiles. In other words, 25% of all WGD tumors possess ploidy below 2.9. Following the reviewer's comment, we have clarified this statement.

Reviewer #3 (Significance (Required)):

The novelty is to look at CNV signatures upon drug treatment (beyind mutational signatures). However, as mentioned above, it is not clear how different exposures that ultimately cause DNA break would have distinct CNV pattern. Overall, the results seem modest to me. Although statistically significant, the increase in specific classes of CNVs in treated v untreated WGD mets is modest (Fig 2d), casting a doubt on clinical significance. This work could still be of interest to some researchers, in particular, those interested in mutational signatures of environmental exposure. This work should be interpreted in the context of pan-cancer signatures of CNVs very recently published https://www.nature.com/articles/s41586-022-04738-6.pdf.

The increase of chromosomal fragments below 10 Mb across platinum-exposed tumors is between 13% and 387% with respect to unexposed tumors (Fig. 4a). The statistical significance of the signal of platinum exposure on the number of CN fragments is smaller than that observed for single nucleotide variants produced by the exposure to platinum (https://www.nature.com/articles/s41588-019-0525-5). However, while each single nucleotide variant affects a single nucleotide, a chromosomal fragment in the middle of the range observed would affect thousands of base pairs. In other words the cumulative effect of the platinum CN footprint on exposed tumors and normal cells is much larger than that of single nucleotide variants. (See also response to point 1.)

With respect to CN signatures, motivated by the reviewer's comments we now demonstrate, using the activity of CN signatures extracted from the HMF cohort (using the methodology presented in https://www.nature.com/articles/s41586-022-04738-6) that none of them is significantly different between tumors exposed or unexposed to major anticancer therapies. Note are there any significant differences in the activities of the original CN signatures extracted in the aforementioned paper across primary tumors between exposed and unexposed tumors in the HMF cohort.

My background is in computational biology, working on transcriptional regulation for decades and more recently in cancer systems biology. I am comfortable with the techniques employed in this work but not so much with the mechanisms linking a specific drug to specific copy number signatures and also with the clinical significance of this problem. Keywords: "Computational biology", "Bioinformatics", "Transcriptional regulation", "NGS", "Omics", "Cancer systems biology"

Reviewer #4 (Evidence, reproducibility and clarity (Required)):

Summary:
This paper by Gonzalez et al attempts to identify copy number footprints of anti-cancer therapies. It follows previous work by the group looking at single base mutational footprints of anti-cancer therapies, which provides clear evidence of the effect of these drugs on the genome. This study into copy number footprints is less convincing, mainly due to the challenges in identifying these low frequency copy number signatures. The authors present weak evidence using CN signatures for an increase in the number of chromosomal fragments less than 10 Mb in size. However, this is not consistently significant between different cancer types treated with the same drugs. The interesting finding of an effect of platinum treatment intensity on copy number is seen quite nicely in Fig 4b when pooled into one simple effect. Signature analysis in this case seems unnecessary as the main finding is that platinum treatment results in increased 10 kb- 10 Mb fragments but only when pooled in this way. The paper is otherwise nicely written, although some clarity adjustments are required in the Figures and Figure legends.

We thank the reviewer for their appreciative summarization of our work.

Major comments:
Whilst a commendable effort has been made to identify copy number footprints, the evidence presented here for the identification of CN signatures is not so convincing. The main focus of the paper is the effect of Platinum based therapies and yet the two featured cancer types lung non-small cell and colorectal do not have consistent significant effects in the signature analysis.

The reviewer is correct that we don't identify a CN signature (in the sense understood in recently published manuscript by Steele et al. https://www.nature.com/articles/s41586-022-04738-6) associated with platinum treatment. A clearer statement to this effect has now been added to the manuscript as a result of novel analyses of these CN signatures in the cohort studied in our work.

What we identify (as the reviewer states in their summary of our work) is a general increase of chromosomal fragments below 10Mb among platinum-exposed tumors. This is consistent across tumor types as shown in Figure 4a, and it is what we describe in the manuscript as the platinum CN footprint. We precisely avoid the term signature in an effort to prevent confusion with the canonical usage of this term.

The referenced BioRXiv paper by Steele et al. is now published https://www.nature.com/articles/s41586-022-04738-6 and one wonders whether additional methods and analyses performed during their peer review may be useful in this paper as well. Can reanalyse using the predefined 21 CN signatures from Steele et al?

We thank the reviewer for this suggestion. Actually, taking advantage of the fact that the tool employed to extract CN signatures de novo (the original SigProfiler aimed at mutational signatures extended to CN features) was available prior to the publication of this article, we had already carried out a CN signatures extraction de novo from the HMF cohort. We then asked if any of these CN signatures (their activity across tumors) is significantly associated with the treatments in the cohort, and found none (current Fig. S3a). In the manuscript we hypothesize that this is due to the intrinsic difficulties in defining CN signatures, as opposed to SBS and DBS signatures. This is why we decided in our study to focus on a collection of individual CN features that show differences between platinum-exposed and unexposed tumors to define the platinum CN footprint.

Following the suggestion by this (and other) reviewer, we have now carried out the same analysis (identification of CN signatures potentially related to exposure to anticancer therapies) using the set of CN signatures originally defined by Steele et al in their paper (reference 21 in our manuscript). This analysis yields negative results too (current Fig. S3c). We also now include the equivalence (established through linear reconstruction) between the CN signatures extracted de novo from the HMF cohort and the CN signatures originally defined by Steele et al. across primary tumors. (This equivalence is provided by the SigProfiler upon extraction.) In general, the signatures extracted across HMF tumors in general bear little resemblance to those extracted from primary tumors (highest cosine similarity of a linearly reconstructed signature, 0.775). This is presented in current Figure S3b.

Taken together, these results further strengthen our point that a CN footprint defined from differences in individual CN features are probably more appropriate than CN signatures in their current format to identify the effects of anticancer therapies on the CN landscape of exposed cells.

From Fig 2a there are 5 cancer types in HMF with Platinum treatment: Lung non-small cell, colorectal, esophagus, urothelial and ovary and it is not clear if all these cancer types are combined or separated in the final analysis. All but urothelial are featured at some point though, but e.g. ovary has no significant differences between treated and untreated. Are samples from all 5 cancer types combined in the "treated versus untreated" analyses?

We thank the reviewer for pointing this out. The analysis is carried out separately by cancer type. We have now included a statement in the **Profiles of chromosomal fragments associated with anticancer therapies** section clarifying this.

The strongest evidence for a real effect on copy number for platinum treatment comes in Figure 4, where there is a significant increase in LoH segments CN 1-4 with samples showing high SBS 35 mutations (a clever idea!). Attempting to separate out the samples into a "Copy number signature" in Figures 2 and 3 seem a bit like fillers to get to this actual potentially interesting finding. What is the benefit of separating out the copy number and zygosity when the real effect is much clearer when you pool everything and simplify it?

As the reviewer points out, and we highlight above, it is this type of chromosomal fragments that we conceptualize as the platinum CN footprint. This, however, is a discovery that stems from the unbiased analysis carried out across tumor types and treatments, the results of which are presented in Figure 2 and which is further characterized in Figure 3. It would have been impossible to identify this footprint from the outset. We reasoned that the CN features defined by Steele et al. were a good starting point to capture differences in the overall landscape of chromosomal fragments of tumors exposed or unexposed to DNA damaging drugs.

Can you investigate other drug treatments using this bulk approach using the proxy of the SBS drug mutations to indicate the "strength" of the mutational process of the drug.

In theory this is possible for treatments that leave both discernible mutational and CN footprints. So far, only platinum-based drugs fulfill this criteria. In the case of 5-FU a salient mutational footprint is associated with the exposure to the drug, but we were unable to identify any discernible CN footprint.

Another point of interest is the overlap between treatments. A judgement call is made as to which is the overriding drug corresponding to the effect. Is it possible to separate these effects with NMF as per SBS? Or could a combination effect be detected? Likely the numbers would be too low for this separated analysis. But just looking at LoH 10kb to 10 Mb might show something?

This is a very interesting suggestion. Several lines of evidence support the idea that the observed CN footprint is associated with exposure to platinum and not 5-FU and that it is not a combination of both drugs (see above response to point 7 raised by reviewer 3). The most important is that the CN footprint is also observed across tumors that are not exposed to 5-FU. For example, in the case of the ovarian tumor cohort, which are not exposed to 5-FU the CN footprint is recovered (Fig. 3c; 4a), although the individual CN features are not significant, due to the low numbers.

With respect to the overlap specifically between platinum-based therapies and pyrimidine analogs, we have included in the manuscript a new Supplementary Table (Table S4) presenting the numbers of WGD tumors exposed to different pairs of drugs across cancer types. We have also extended the statements about the overlaps in the manuscript to further clarify the decisions made. (See above our reply to point 7 by reviewer 3.)

A table summarizing the included samples, treatments, overlap, etc per cancer type is missing.

Following the reviewer's suggestion, we have prepared and included in the manuscript two new Supplementary Tables. Table S2 details the number of WGD and non-WGD metastatic tumors of each tumor type across the HMF cohort exposed to different anti-cancer therapies. Table S4 presents the number of tumors exposed to different pairs of treatments across tumor types in the cohort.

Your study may also benefit from a comparison to the latest Hartwig cohort paper https://www.biorxiv.org/content/10.1101/2022.06.17.496528v1.full particularly focusing on the suggestion of Treatment enriched drivers (TED) some of which are infact copy number driven.

We thank the reviewer for this interesting suggestion. The only TED identified by the authors, which is related to platinum-based drugs (with the criteria described in the Supplementary Table 8 of their manuscript) concerns point mutations of TP53 across metastatic stomach adenocarcinomas (where we are unable to identify the CN footprint due to a low sample size). Although some driver amplification or deletion events do appear significantly enriched across platinum exposed tumors of different cancer types, they are discarded by the authors due to lack of orthogonal evidence of being associated with the specific mechanism of action of the drug.

We now include this observation in the revised manuscript:

*As anticipated, we observed that chromosomal fragments smaller than 10 Mb (representative of the platinum CN signature) are evenly distributed along the genomes of WGD colorectal and lung tumors (Fig. 3b; Fig. S4a,b). Had this increase in the number of platinum-related chromosomal fragments across exposed tumors been constrained to one or few genomic regions, it would point to positive selection of one or more resistance-associated genes. A recent systematic analysis of the HMF cohort revealed that only mutations in TP53 across stomach adenocarcinomas appear as a potential bona fide driver event associated with the exposure to platinum in the HMF cohort (Martínez-Jiménez et al, 2022).*

Minor comments:
The Figure legends and Figures themselves need to be altered for clarity. Axis should be labelled more specifically, e.g Fig 1d axis currently reads "percentage". Fig 3b says left and right and there is no such thing. What do the sizes of the circles in 2c represent? Can you indicate cancer type as well in this plot (shading or line type) or are all treated samples pooled- this is not clear?

We thank the reviewer for this suggestion. We have checked all figures and figure legends to enhance their clarity.

It is not clear why Fig 2a only includes HMF samples and not PCAWG- PCAWG could be in supplement?

Figure 2a describes the types of anticancer treatments received by patients bearing different types of malignancies. PCAWG tumors are primary and treatment-naive; this is why all the study to identify treatment-related CN features focuses on the HMF cohort.

Be consistent with labelling as well, in the text everything is referred to as 10 kb -10 Mb and some Figures labeled as such but others with 10^4- 10^7. How is the size 10 kb established? All the plots show 0-100 kb, where did the 10 kb limit come from?

We thank the reviewer for this recommendation. It actually led us to review our definition of the platinum CN footprint and to realize that, indeed, fragments smaller than 10 kb are part of this footprint. All analyses (and relevant figures and supplementary tables) have been updated accordingly. The rationale to define the CN footprint is now more thoroughly explained (Fig. 3a).

Why is ovary not in 4b?

There are very few platinum-exposed ovarian tumors with WGD and activity of SBS35. Therefore, the groups of tumors with high and low activity of SBS35 are too small to carry out a meaningful comparison of the platinum CN footprint in Fig. 4b.

Methods needs clarification. Are the visualized samples the average of the cancer types in each of the two groups (untreated vs treated) how many samples in each group? The table suggested above would help a lot with understanding what is actually being compared. How reliable is the calculation of WGD status? Some explanation into the values used in the calculation "WGD: 2.9 -1.7*LoH <= Ploidy" is warranted.

Following the reviewer's suggestion, we have added two new Supplementary Tables (Tables S2 and S4) presenting a more thorough description of the subset of HMF tumors employed in the detection of treatment-related CN features (Table S2) and the overlap between treatments in terms of numbers co-treated tumors (Table S4). We have also expanded the rationale behind the inequality used to separate WGD and non-WGD tumors (see new version of Methods).

CROSS-CONSULTATION COMMENTS
Everyone seems to be in relative agreement that the results are modest, should be compared to https://www.nature.com/articles/s41586-022-04738-6.pdf and require some clarity throughout the manuscript.
Additional analyses, such as comparing to other datasets such as POG570 would benefit the paper.

Reviewer #4 (Significance (Required)):

Whilst previous studies have looked at the effect of anti-cancer drugs on single base mutations, owing to the challenges also seen here, no thorough investigation of the effect of these drugs on copy number has been performed. Therefore, this is an advance, albeit minor as the "copy number signature" of the exposed cancers was not particularly clear.
The use of WGD samples was a clever step forward for the analysis of copy number, as the effects of selection are weakened with an extra copy of the genome.
The finding that increased treatment with platinum results in increased copy number changes of size 10 kb to 10 Mb is an interesting finding, and something that could be considered when looking at treatment options in the future, particularly if it is shown to also affect normal cells in this way.
The cancer genomics field in which I am a part, would be interested in this finding.

We thank the reviewer for their appreciative comment of our work.
